# Taxonomy of introns, their evolution, and the role of minor introns in stress response

**DOI:** 10.1101/2022.10.12.511939

**Authors:** Anouk M Olthof, Charles F Schwoerer, Audrey L Weber, Iswarya Arokiadhas, Karen Doggett, Stephen Mieruszynski, Avner Cnaani, Joan K Heath, Jakob Biran, Rahul N Kanadia

## Abstract

Despite the high conservation of minor introns across eukaryotic supergroups, specific lineages have completely lost minor intron splicing, which has raised questions about their evolution and purpose. Addressing these questions requires identification of the introns that are affected by minor spliceosome inhibition. To this end, we applied principles of Linnaean taxonomy combined with position-weight matrices to produce five intron classes: minor, minor-like, hybrid, major-like and major. We classified introns across the genomes of 263 species of six eukaryotic supergroups, which can be viewed at the Minor Intron Database (MIDB). Transcriptomic analysis revealed that ∼40% of the minor introns are responsive to minor spliceosome inhibition, while an additional 5% of the minor-like and hybrid introns are also affected. We propose that minor-like introns represent an intermediate in the conversion of minor to major introns and uncover the importance of a guanine at the −1 position of the 5’ splice site in facilitating this shift in spliceosome dependence. Finally, we find that minor introns are aberrantly spliced in fish and plants upon cold stress, thereby providing a potential explanation for their high degree of conservation in these lineages.

## Introduction

One of the major hallmarks of eukaryotic evolution has been the emergence of spliceosomal introns through invasion of self-splicing group II introns into the genome. Indeed, analysis of orthologous genes suggests that the last eukaryotic common ancestor (LECA) and ancestors of each eukaryotic supergroup had intron-rich genes (1, 2). Even though almost all eukaryotic genomes still contain introns, the intron density between lineages varies greatly, suggesting that different species have either lost or gained introns. The evolution of most eukaryotic lineages is thought to have involved the substantial loss of introns, exemplified by the presence of very few introns in many unicellular organisms (1). At the same time, intron gain is thought to have accompanied major eukaryotic transitions, such that metazoa generally possess intron-rich genomes (1, 2). The presence of introns in eukaryotic genomes resulted in the need for a machinery capable of their removal, so that genes could be properly expressed. Indeed, the spliceosome, consisting of five small nuclear RNAs (snRNAs) and many associated proteins, is believed to have co-evolved with spliceosomal introns (3). Specifically, the snRNAs are thought to have originated from the catalytic fragments of group II introns and are highly conserved across eukaryotic lineages (3–5).

The splicing of introns is accomplished through base-pairing of spliceosomal snRNAs with highly conserved sequences at the 5’ splice site (SS), branch point sequence (BPS) and 3’SS of the intron. The identification of two distinct spliceosomes, each consisting of different snRNAs but sharing many of the associated proteins, led to a binary classification of introns (6–9). Most introns possess consensus sequences that are complementary to the components of the major (U2-type) spliceosome and are therefore referred to as major introns. In contrast, a small subset of introns with divergent consensus sequences, called minor introns, are recognized by the minor (U12-type) spliceosome. While binary intron classification based on splice site sequence has proven useful for elucidating the mechanism of minor intron splicing, next generation sequencing of genomes and transcriptomes has revealed a wide diversity in consensus sequences. Moreover, several lines of research have reported that not all bioinformatically classified minor introns are affected upon minor spliceosome inhibition (10–14), suggesting that this dichotomous classification of introns is an oversimplification. Gaining insight into the targets of each spliceosome will enhance our ability to uncover the mechanism of disease pathogenesis of spliceosomopathies, a group of inherited disorders caused by mutations in spliceosome components (15). In the past decade, there has been a vast increase in the number of annotated genomes that are publicly available, facilitating a revision of intron classification across a diverse group of species.

Classification of introns across a diverse group of species may shed light on the evolution of minor introns. Both minor spliceosome components and minor introns have been identified in many eukaryotic lineages, although they are reportedly absent in species such as yeast, green algae, *Dictyostelium*, and *Caenorhabditis* (5, 16, 17). This apparent paradox of high conservation across eukaryotic supergroups, but complete loss of minor intron splicing in other lineages has raised questions about the origin and evolution of minor introns. Minor introns, while likely ancient and present in LECA, are thought to have emerged after major introns (17, 18). The eukaryotic species lacking minor introns and/or minor spliceosome components could have lost minor introns either through homologous recombination of a reverse transcribed spliced mRNA with the genome, or through conversion of the minor intron to a major intron (18, 19). The latter would be relatively easy, as only a few sequential point mutations would be required to obtain major-type consensus sequences (20). If conversion of minor introns to major introns occurs, we would expect to detect introns that are in flux. These introns would possess degenerating consensus sequences that might not be recognizable by either the major or the minor spliceosome. Additionally, introns with a minor-type 5’SS and major-type 3’SS, or vice versa, might also exist. In fact, a few so-called hybrid introns have previously been identified using position-weight matrices (PWMs) (21). However, since the classification of introns has predominately focused on opisthokonta, a good understanding of minor intron evolution across different eukaryotic lineages remains enigmatic.

Ultimately, gaining insight into the evolution of minor intron splicing might be able to provide us with an explanation of an exciting, yet unanswered question: how have evolutionary pressures ensured the maintenance of minor introns in a subset of genes? Here we applied a non-binary classification of introns based on consensus sequences across 263 genomes from all eukaryotic supergroups, which can be accessed at http://midb.pnb.uconn.edu. By leveraging transcriptomic data, we identified that ∼40% of all minor introns and a small subset of minor-like and hybrid introns are aberrantly spliced when the minor spliceosome is inhibited. We found that these minor-like introns, whose consensus sequences resemble that of minor introns, except for a guanine at the −1 position of the 5’SS, represent a transition in the conversion of minor introns to major introns. While this reveals a mechanism by which some minor introns are lost during evolution, the majority of minor introns are highly conserved. Importantly, the most highly conserved minor introns are found in genes that participate in the stress response, providing a potential explanation for their maintenance. Indeed, cold stress in both fish and plants results in elevated minor intron retention, suggesting that the minor spliceosome and minor introns are under active selection by one of the major drivers of evolution, changes in temperature.

## Materials and Methods

### Classification of introns

The classification of introns was done using PWMs, with minor changes to previously described methods (22). The genome and intron data for all 263 species was extracted from FASTA and GTF files obtained from Ensembl **(Table S1)**. Introns were initially binned as putative major (GT-AG and GC-AG), putative minor (AT-AC), or ‘other’ based on their terminal dinucleotide sequence. For putative major introns, an initial PWM for the 5’SS was generated from the −2 to +6 nucleotides, while for putative minor introns, an initial 5’SS PWM was constructed using the +4 to +9 nucleotides. These nucleotide sequences were selected due to their importance for the base-pairing with U1/U6 and U11/U6atac, respectively (23). The initial major PPT PWM was then constructed from the −13 to −1 nucleotides of putative major introns. For the construction of an initial minor BPS PWM, a sliding window from the −40 to −1 position was first applied to all putative minor introns to extract all potential twelve nucleotide BPS with an adenine at the +9 or +10 position. This initial PWM was then utilized to score all putative BPS generated by the sliding window. The highest scoring BPS from each putative minor intron with a positive cumulative LOD score was extracted to generate two initial minor BPS PWM: one with adenine at the +9 position, and one with adenine at the +10 position. After generation of the initial PWMs **(Fig. S1)**, all introns were scored and re-binned as described in **Fig S2A** to generate refined PWMs for the major and minor 5’SS, minor BPS and major PPT. After rescoring all introns against these refined PWMs, introns were binned using the scoring criteria outlined in **Fig. S2B**. This intron classification for all 263 species is available at the Minor Intron Database, which can be accessed at http://midb.pnb.uconn.edu (22).

### RNAseq of rnpc3 mutant zebrafish

RNA from two independent *rnpc3* mutant alleles and their respective wildtype controls (n=3 for each genotype) were sequenced. *Clbn*^*s846*^ identified in an ethylynitrosourea mutagenesis screen encodes a T to A transition in intron 13, creating a novel 3’SS 10 nucleotides upstream of the canonical 3’SS (12). This results in aberrant transcripts all containing premature stop codons with no correctly spliced exon 13-14 junction transcript detected. *Clbn*^*ZM*^ harbors a retroviral insertion in intron 1 of *rnpc3*; both alleles are functionally null (12). Total RNA was extracted from pools of genotyped 72-hour post fertilization (hpf) larvae using TRIzol. Poly(A)-enriched RNA was used to generate cDNA libraries and sequenced using the Illumina HiSeq by the Australian Genomics Research Facility (AGRF), yielding 100bp single end reads.

### Identification of responsive introns

All introns classified as minor, minor-like or hybrid were used to generate BED files for splicing analysis. For the analysis of major and major-like intron splicing in cold stress conditions, a randomized list of 1000 introns was generated. For introns found in multiple transcripts, the longest transcript was considered as canonical, and used to extract the coordinates of the flanking exons. A list of the RNAseq datasets that were used in this study can be found in **Table S2**. Reads were aligned to the Ensembl v.99 genome assembly of respective species using Hisat2 (24). Retention and alternative splicing analysis were then performed as described previously (11, 22). Significant differences in mis-splicing indices were determined by welch’s t-test or one-way ANOVA, followed by post-hoc Tukey test using custom R scripts.

### Identification of minor spliceosome snRNAs

All available sequences for U11 (RF00548), U12 (RF00007), U4atac (RF00618) and U6atac (RF00619) were downloaded from the Rfam database. For species that did not have any minor spliceosome snRNA recorded in the Rfam database, a blastn database was built from the genome FASTA file. We then performed blastn queries (word size=7) on the Rfam fasta file with annotated snRNA sequences against these databases to identify putative snRNA homologs in other species. All coordinates of hits with E value < 10 were extracted and the sequences + flanking 200 nucleotides were identified. These putative snRNA homolog sequences (solely based on sequence similarity) were further analyzed using the Infernal package (25). To this end, the covariance models for U11, U12, U4atac and U6atac were downloaded from the Rfam database. This model was then run against the putative snRNA sequence fasta files using cmsearch. To identify high-confidence snRNA homologs, we first established score thresholds by comparing the covariance model for each snRNA that was downloaded from the Rfam database to the annotated snRNA sequences also downloaded from the Rfam database. The score corresponding to the lowest scoring hit within the inclusion threshold was considered the threshold for identifying new snRNAs. These were: 37.3 for U11, 53.0 for U12, 53.2 for U4atac, and 22.9 for U6atac. All putative snRNA homologs above these thresholds were therefore considered reliable hits. If multiple hits were above this threshold, these could be gene copies or pseudogenes; if no hits were above this threshold but there were hits above the inclusion threshold set by cmsearch, they were analyzed separately.

### Intron orthology

DIAMOND protein databases were generated for each species using the protein FASTA files obtained from Ensembl and custom scripts. Proteins were blasted against each of the protein databases using DIAMOND with the following options: –min-orf 1 and –e 10E-10. For each intron, the canonical transcript was defined as the longest protein-coding exon sequence that contained the intron. If an intron was found outside of the ORF of the canonical transcript, the next longest sequence in which the intron was located within the ORF was extracted. Orthologous genes were then identified by blasting the translated sequence of the canonical transcript against all DIAMOND databases. To identify orthologous introns, we pairwise aligned the sequence of the ten amino acids flanking all exon-intron boundaries of orthologous genes to the flanking ten residues of the reference intron using the pairwise2 package from Biopython. Introns with the highest BLOSUM-62 score were then extracted (based on highest match score if BLOSUM-62 was tied) and considered orthologous if the conserved amino acid identity was at least 40%.

### Phylogeny

Phylogeny of species was downloaded from the NCBI Taxonomy browser, and modified using the criteria described in (26). Phylogenetic tree with branch lengths reflecting time was obtained from timetree.org.

### RT-PCR analysis on cold-exposed Nile tilapia

The experiments were approved by the Agricultural Research Organization Committee for Ethics in Using Experimental Animals. For the chronic cold stress experiments, healthy male *Oreochromis niloticus* were either maintained at 27 ºC (control) or at 14 ºC (cold exposed) for three weeks. Midbrains were then dissected and total RNA was extracted using TRIzol according to the manufacturer’s instructions. cDNA was prepared from DNase-treated RNA and used for PCR analysis. Primer sequences used to detect intron retention can be found in **Table S3**. All spliced products were excised and confirmed using Sanger sequencing.

## Results

### Identification of introns responsive to minor spliceosome inhibition

Inherent to the classification of a minor intron is the assumption that its splicing will be disrupted upon minor spliceosome inhibition. However, ubiquitously expressed minor intron-containing genes (MIGs) are spliced in a cell type specific manner (22). This variation in the responsive nature of minor introns complicates the construction of a comprehensive list of introns that are dependent on the minor spliceosome. Nevertheless, development of such a list is essential for benchmarking bioinformatics pipelines aimed at annotating introns and for identifying the biological pathways regulated by the minor spliceosome, as the minor spliceosome has recently been suggested as a potential therapeutic target for cancer (27, 28). We therefore decided to analyze all currently available transcriptomic datasets in which the minor spliceosome was inhibited **(Table S2)**, yielding a comprehensive list of experimentally validated, “responsive” introns in the five model organisms: human (*Homo sapiens*), mouse (*Mus musculus*), zebrafish (*Danio rerio*), fruit fly (*Drosophila melanogaster*) and maize (*Zea mays)*. To increase our resolution, we first made modifications to our previously published classification pipeline such that introns were no longer binned into either minor or major class (22). Rather, they are grouped into one of six categories on a spectrum bookended by minor and major introns, including: minor introns, minor-like introns, hybrid introns (consisting of a major-type 5’SS and minor-type BPS or vice versa), major-like introns, major introns and non-canonical introns. While the PWM score for the 5’SS sequences was highly informative for the classification of introns, the score for the BPS/PPT was less discriminatory **(Fig. 1A)**. In all, we identified 845, 711, 659, 18 and 427 minor introns in the genomes of the human, mouse, zebrafish, fruit fly and maize, respectively **(Fig. 1B)**. These numbers are comparable to previous reports (**Fig. S3**) (21, 29, 30). Moreover, we found a similarly sized population consisting of minor-like introns and hybrid introns to that of minor introns in these model organisms **(Fig. 1B-C)**. Closer examination of the consensus sequences for each intron class in the human genome revealed that non-canonical introns do not have a distinct motif at either the 5’SS or the 3’SS, in agreement with their LOD scores being below 0 (see methods) **(Fig. 1A,C)**. In contrast, the consensus sequences of major-like introns are like those of major introns, although major-like introns generally possess a weaker PPT. Similarly, the consensus sequences of minor-like introns resemble those of minor introns. The main difference is a significantly reduced conservation of the C at the +5 and +6 positions of the BPS in minor-like introns, as well as a significant enrichment of a G at both the −1 position of the 5’SS and the +1 position of the 3’SS (*P*<0.0001; Chi-squared test). Similar patterns were observed for introns in the mouse, zebrafish, and maize genomes **(Fig. S4)**. Notably, the G at the −1 position of the 5’SS is a hallmark feature of major introns and is normally used by U1 snRNA for base-pairing (31). The presence of this nucleotide in minor-like introns therefore raises the possibility that these introns can be recognized by the major spliceosome, despite the overall resemblance of their consensus sequences to minor introns.

**Figure 1.**
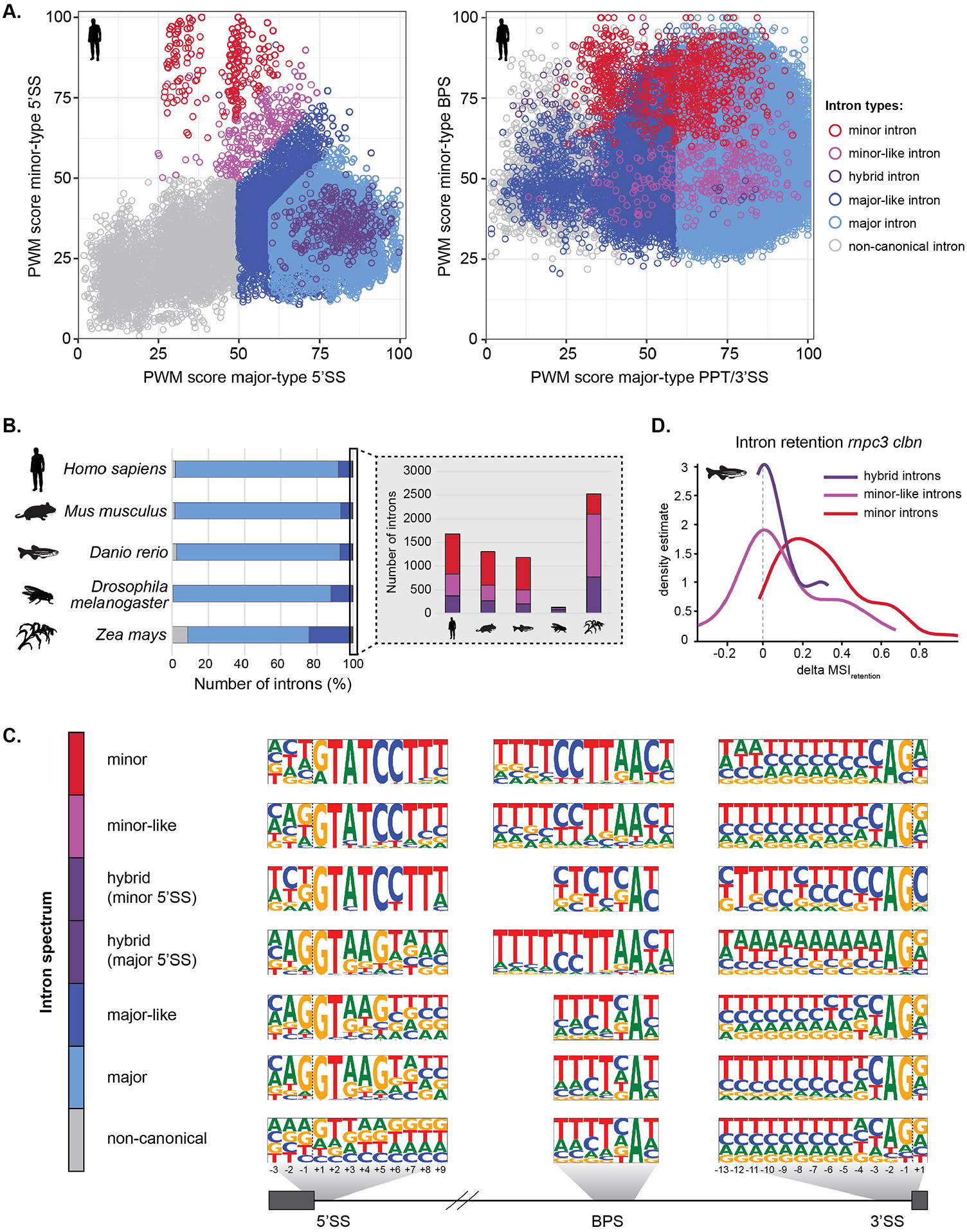
Classification of introns in model organisms. **(A)** Scatterplot with PWM scores for all human introns. Color-coding indicates intron class. **(B)** Stacked bargraph for the distribution of intron classes in *Homo sapiens, Mus musculus, Danio rerio, Drosophila melanogaster, Zea mays*, with zoomed inset showing the number of minor, minor-like and hybrid introns. **(C)** Frequency logos of consensus sequences for 5’SS, BPS and 3’SS of the different intron classes in the human genome. Dashed line denotes the exon-intron boundary. **(D)** Density plot for the level of retention of minor, minor-like and hybrid introns in the rnpc3 clbn compared to wild-type zebrafish. MSI=mis-splicing index.

Next, we studied intron retention and other *de novo* alternative splicing events around introns in twenty publicly available RNAseq datasets that covered various cell types in human, mouse, fruit fly and maize. Additionally, we performed RNAseq of zebrafish larvae with two different mutations in *rnpc3*, a critical minor spliceosome-specific protein (12). As expected, we found that bioinformatically classified minor introns were retained at higher levels upon minor spliceosome inhibition, reflected by a positive difference in mis-splicing index between the different experimental and their respective control conditions **(Fig. S5-S9)**. However, we observed large differences in the degree of minor intron mis-splicing between datasets, underscoring the importance of integrating multiple datasets for the identification of true minor spliceosome targets **(Fig S5-S9)**. The intersection of introns whose splicing was significantly altered upon minor spliceosome inhibition in at least one dataset revealed that approximately 40% of all minor introns in the human genome were “responsive” in at least one cell type (**Fig 2A**). The splicing of an additional 5% of human minor-like and hybrid introns was also affected by minor spliceosome inhibition **(Fig. 2A)**. Similar numbers of responsive introns were observed for mouse, zebrafish, fruit fly and maize. Even though we anticipate identification of an increased number of responsive minor introns with the generation of more knockout models and deeper transcriptomic data, the current list provides a critical insight into the introns that are highly dependent on minor spliceosome function for their splicing.

**Figure 2.**
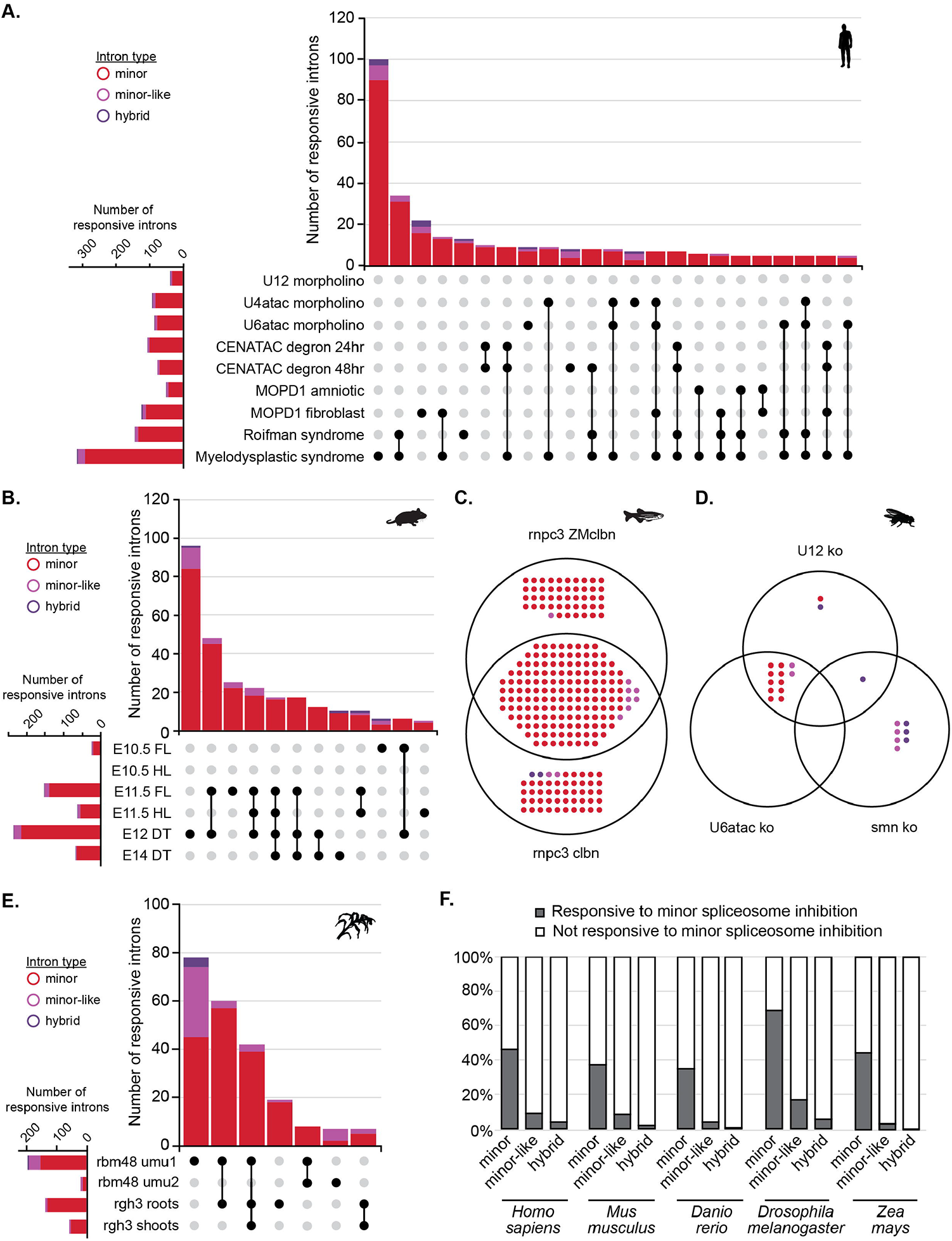
Identification of introns responsive to minor spliceosome inhibition. **(A-B)** Upset plot for mis-spliced introns in different **(A)** human and **(B)** mouse datasets in which the minor spliceosome is inhibited. **(C-D)** Venn diagram for mis-spliced introns in different **(C)** zebrafish and **(D)** fruit fly datasets in which the minor spliceosome is inhibited. **(E)** Upset plot for mis-spliced introns in different maize datasets in which the minor spliceosome is inhibited. **(F)** Bargraphs with total number of responsive minor, minor-like and hybrid introns in the different model organisms. For more information on the RNAseq datasets used, see also Table S2.

### The presence of a G at the −1 position of the 5’SS reduces the dependence of minor-like intron on the minor spliceosome

We next wanted to gain insight as to why certain introns are more affected by minor spliceosome inhibition than others. We did not observe a significant difference in the consensus sequences at each splice site between responsive and unresponsive minor introns **(Fig. 3A)**. Surprisingly, we did note a difference in the 5’ SS consensus sequence of responsive and unresponsive minor-like introns in humans **(Fig. 3A)**. Specifically, there was a bias against the G at the −1 position for minor-like introns affected by minor spliceosome inhibition. Similarly, we noted a bias against the G at the +1 position in the 3’SS of responsive minor-like introns **(Fig. 3A)**. The close resemblance of consensus sequences of responsive minor-like introns to that of minor introns suggests that the presence of a guanine at the exon-intron boundaries reduces the reliance on the minor spliceosome and might allow recognition of these introns by the major spliceosome. In addition to a difference in consensus sequence, we found that the distance of the BPS from the 3’ end of responsive minor-like introns was significantly shorter than that of unresponsive minor-like introns **(Fig. 3B)**. The distribution of these BPS distances also resembled that of minor introns, while the distribution for unresponsive minor-like introns was more uniform **(Fig. 3B; Fig. S10)**.

**Figure 3.**
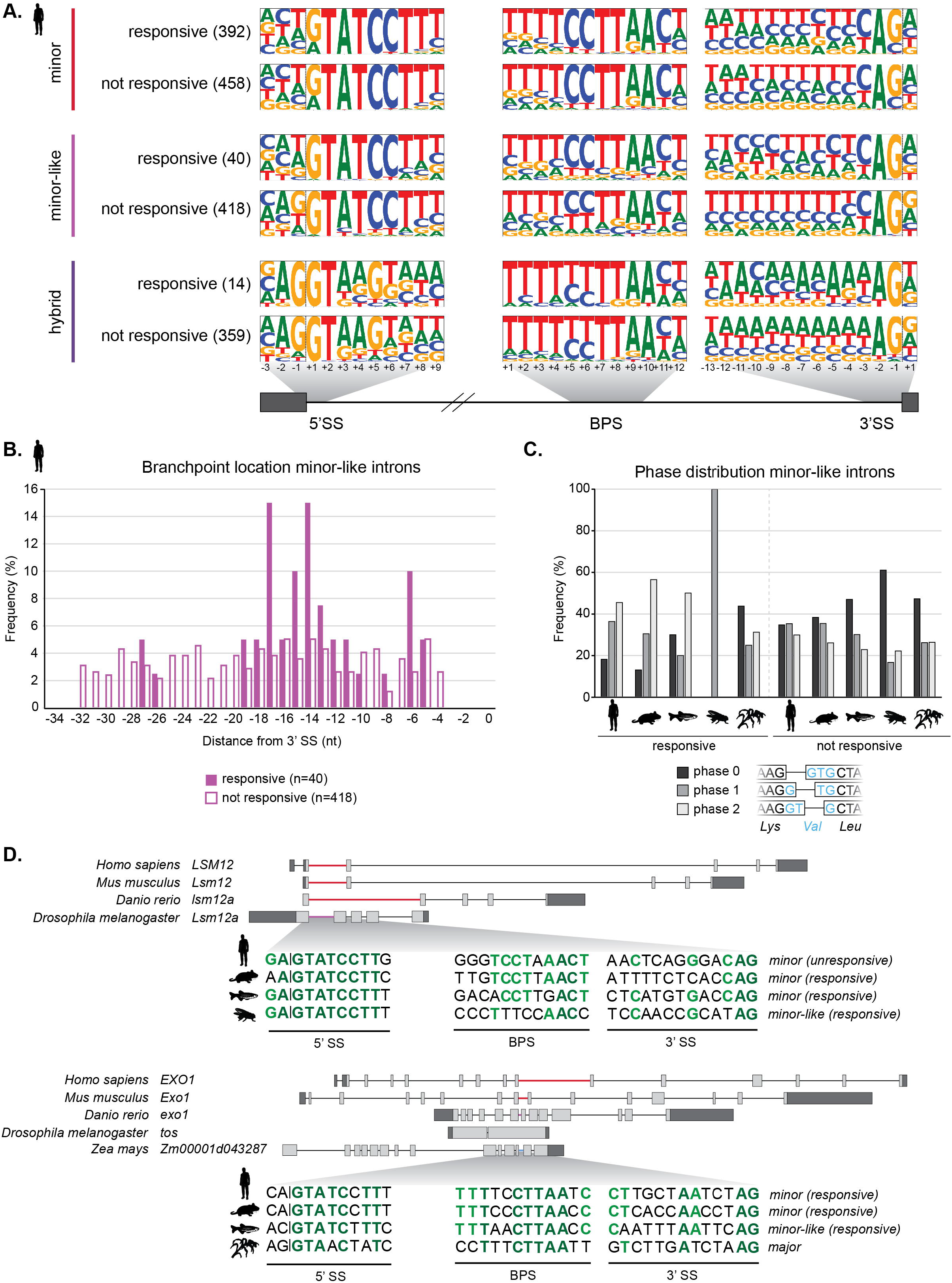
Features of responsive minor-like introns resemble that of minor introns. **(A)** Frequency logos of consensus sequences for 5’SS, BPS and 3’SS of human minor, minor-like and hybrid introns whose splicing is (responsive) and is not (unresponsive) affected by minor spliceosome inhibition. Number in brackets denotes the number of introns used in each group to create frequency logos. Dashed line denotes the exon-intron boundary. **(B)** Histogram of the distance of branchpoint adenosine from the 3’ exon-intron boundary for responsive and unresponsive minor-like introns in the human genome. **(C)** Phase distribution for responsive (left) and unresponsive (right) minor-like introns in model organisms. **(D)** Gene schematics and splice site sequences for orthologous intron cluster for LSM12 and EXO1. Color coding of intron represents intron class. Nucleotides in dark green are 100% conserved between orthologous introns, nucleotides in light green are 75% conserved.

Introns are not positioned randomly in the genome; it has long been known that an excess of major introns is found in phase 0 (i.e., positioned between two codons), while minor introns are predominately found in phase 1 (after the first nucleotide of a codon) or 2 (after the second nucleotide of a codon) (18, 32). The reason for this position bias remains unclear, but it has been proposed that minor introns in phase 0 could be more easily converted to a phase 0 major intron (29). Interestingly, we found that unlike major and minor introns, minor-like introns in the human and mouse genome were equally distributed between the phases **(Fig. S11)**, thereby providing support for the idea that minor-like introns resemble an intermediate state in the conversion of minor introns to major introns. If such a conversion were to take place, one might predict that minor-like introns are enriched in orthologous intron clusters containing minor introns. Indeed, we found several examples of orthologous introns containing both minor and minor-like introns. For instance, the minor-like intron in the *Drosophila Lsm12a* gene is orthologous to a minor intron in human, mouse, and zebrafish **(Fig. 3D)**. While the 5’SS sequence is almost entirely identical between these introns, the BPS in *Drosophila* deviates substantially. Similarly, an orthologous intron cluster in *exo1* consists of a minor intron in human and mouse, a minor-like intron in zebrafish, and a major intron in maize **(Fig. 3D)**. In contrast, the orthologous gene in *Drosophila, tos*, appears to have lost this intron altogether. Importantly, the minor-like introns in these orthologous minor intron clusters are often still responsive to minor spliceosome inhibition, despite their suboptimal consensus sequences. In fact, orthologous intron clusters with human responsive minor-like introns contain more minor introns than orthologous intron clusters with unresponsive minor-like introns (Mann-Whitney U test, P<0.0001). In addition, we found that responsive minor-like introns were biased against phase 0, like minor introns, while unresponsive minor-like introns were not **(Fig. 3C; Fig. S12)**. This suggests that intron phase is not only the consequence of an evolutionary process but may also play a role in the splicing reaction.

### Classification of minor, major and hybrid introns across diverse eukaryotic genomes

To further study the conservation and evolution of minor introns, we expanded our intron classification to the genomes of 263 species distributed over six of the seven eukaryotic supergroups defined to date (33). Specifically, we interrogated the genomes of 28 species in the TSAR supergroup, one species in Haptista, three species in Cryptista, 33 genomes in Archaeplastida, 192 genomes in the Amorphea supergroup, and six genomes in the Excavata supergroup **(Fig. S13)**. We identified minor introns in the genomes of all 104 investigated chordates, as well as other opisthokonts such as annelids, arthropods, cnidarians, and mollusks **(Fig. 4; Table S4)**. We also identified minor introns in all streptophytes, including land plants and the green algae charophytes. In all, minor intron density was relatively consistent among chordates and, on average, highest in mollusks (0.26%) **(Fig. 4)**. This was much more variable in arthropods, as the tick *Ixodes scapularis* possessed over 10-fold more minor introns than diptera and many other arachnids **(Fig. 4; Table S4)**. Similarly, the genome of the myxozoan endoparasite *Thelohanellus kitauei* contained far fewer minor introns than *Nematostella vectensis*, another cnidarian, but 5-fold more minor-like and hybrid introns **(Fig. 4; Table S4)**. Surprisingly, even though it has previously been reported that nematodes have lost minor introns, we also identified several minor introns in the genomes of *Caenorhabditis elegans, Loa loa* and *Trichinella spiralis*, among others **(Fig. 4; Table S4)** (17, 21). The correct annotation of the exon-intron boundaries of these genes was confirmed using RNAseq data (data not shown). This discrepancy might in part be explained by the fact that we interrogated all unique introns in the genome, and did not restrict our analysis to the longest isoform for intron classification (21, 29). These minor introns in nematodes made up ∼0.02% of all introns and were therefore 10-fold less abundant than in chordates. In contrast, we found that the percentage of minor-like introns was increased almost 3-fold in nematodes compared to chordates **(Fig. 4)**. This raises the possibility that minor introns are being lost in nematodes through conversion to minor-like introns. Other opisthokonta that have previously been reported to lack minor introns are fungal lineages, most notably the yeast *Saccharomyces cerevisiae*. Consistently, we detected few-to-none minor introns in the *Ascomycota, Basidiomycota* and *Mucoromycota* lineage **(Fig. 4; Table S4)**. The exceptions to this trend were the zoopagomycote *Basidiobolus meristosporus* and neocallimastigomycote *Neocallimastix californiae*, which possess a relatively high minor intron density of 0.5% (237) and 0.2% (129), respectively **(Fig. 4; Table S4)**. Other organisms worth highlighting include the variosean amoebas *Planoprotostelium fungivorum*, which contains a substantially higher number of minor introns (0.14%) than other amoebozoa and *Dictyostelium discoideum*, where more than 5% of all introns were classified as hybrid **(Fig. 4; Table S4)**. Finally, we did not observe any minor introns in red algae or the green algae chlorophytes.

**Figure 4.**
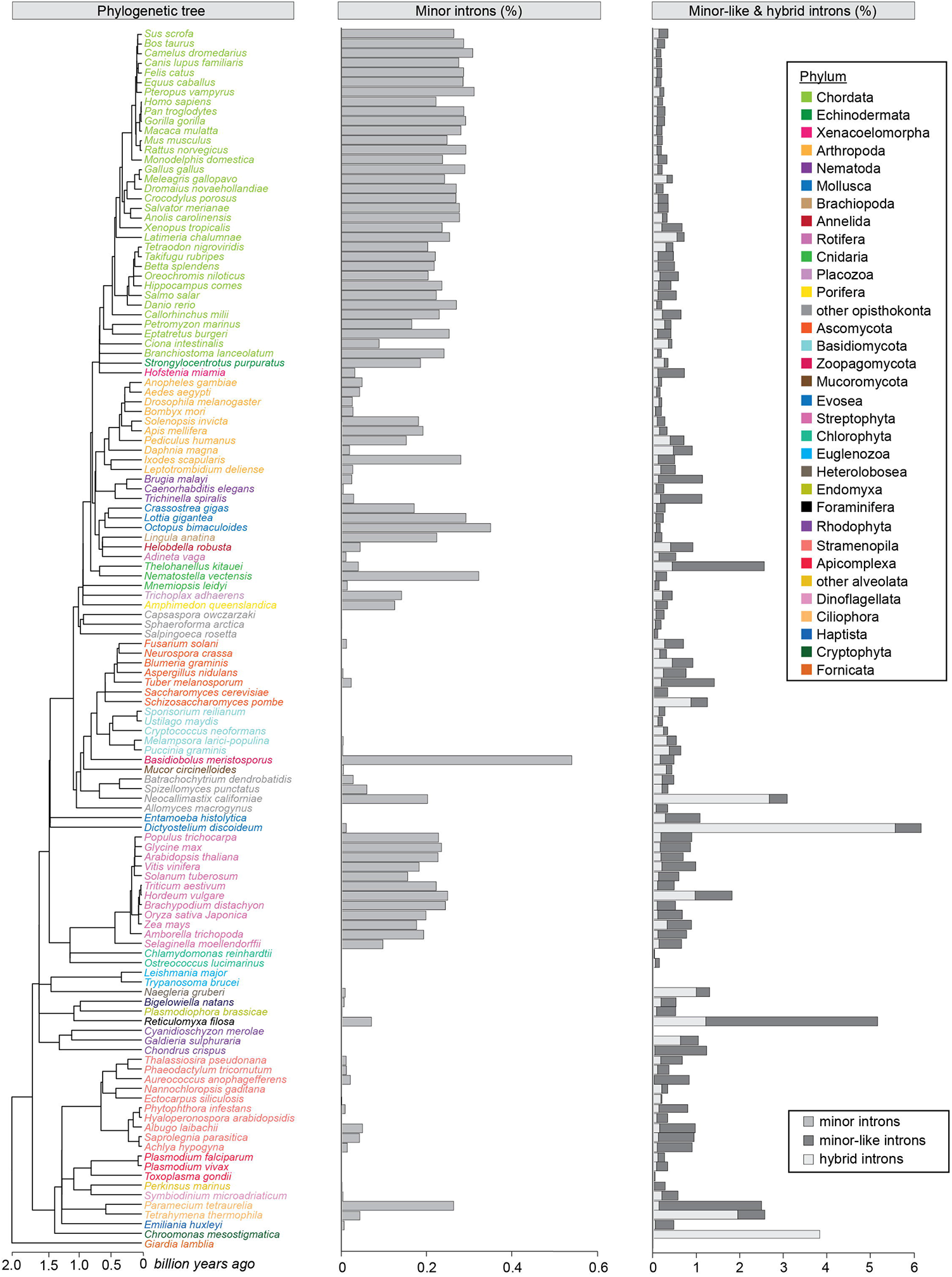
Intron classification in diverse eukaryotic organisms. Phylogenetic tree (left) and bargraph with percentage of minor introns (middle) and minor-like + hybrid introns (right) detected in the genomes of a select set of eukaryotic organisms. Species are color-coded by phylum. For a full list of intron numbers in 263 eukaryotic organisms, see also Table S4.

Relatively uncharacterized so far has been the distribution of minor introns in the more extant Cryptista, Haptista, Excavata and TSAR supergroups. Our analysis revealed the presence of a handful minor introns and a substantial number of minor-like introns in the haptophyte *Emiliania huxleyi* **(Fig. 4; Table S4)**. To our knowledge, this is the first time that minor introns have been identified in Haptista. However, due to the lack of sequenced genomes in this supergroup, it remains unclear whether this is a general feature of this supergroup or specific to this organism. Additionally, we identified minor introns in the genome of the cryptophyte *Guillardia theta*, which is much more intron-rich than that of the other investigated cryptophyta **(Fig. 4; Table S4)**. In contrast, we did not detect strong evidence for minor introns in any of the analyzed species in the Excavata supergroup. Finally, we detected minor and minor-like introns in multiple organisms from the alveolata, rhizaria and stramenopila lineages. Most notably, these include the bygiria *Blastocystis hominis* (0.29% minor, 2.6% minor-like introns) and the foraminifera *Reticulomyxa filosa* (2.1% minor-like) **(Fig. 4; Table S4)**. In all, these findings suggest that minor introns date back approximately 1.7 billion years and are highly conserved across evolution, especially in modern genomes.

### Conservation of minor spliceosome components

Given that minor introns require the minor spliceosome, we next explored the conservation of the unique snRNA components of the minor spliceosome, i.e., U11, U12, U4atac and U6atac (6, 7). The function of snRNAs not only depends on their nucleotide sequence, but also secondary structure. Therefore, we employed blastn in combination with cmsearch to identify putative snRNA genes among all species that we investigated for the presence of minor introns. Using this approach, we identified genes encoding the minor spliceosome snRNAs in almost all metazoans, except for the cnidarian *Thelohanellus kitauei* and most nematodes, though we did detect genes encoding the U11, U12 and U6atac snRNA in the nematode *Trichinella spiralis* **(Fig. 5)**. Similarly, while we did not detect any minor spliceosome snRNAs for species in the fungal clades Ascomycota or Basidiomycota, we did detect several components in the mucoromycota and zoopagomycota lineages **(Fig. 5)**. Moreover, we detected minor spliceosome snRNAs in the amoebozoa *Acantamoeba castellanii* and *Planoprotostelium fungivorum*. In the Archaeplastida supergroup, we identified minor spliceosome snRNAs in land plants but not in green or red algae, which is consistent with the absence of minor introns in these species **(Fig. 4-5)**. Finally, we identified genes encoding U11, U12, and U6atac snRNA in the gyrista clade of stramenopiles, but not in other stramenopile clades such as diatoms, nor in alveolates or rhizaria **(Fig. 5)**. Interestingly, we found that while U4atac snRNA was well conserved among the opistokontha lineage, it was entirely absent in all other supergroups despite the presence of other snRNAs in these lineages **(Fig. 5)**. The only exception to this was *Achlya hypogyna*, a facultative parasite of the stramenopile lineage, where we detected a putative U4atac snRNA **(Fig. 5)**. Normally, the splicing of introns requires a high number of assembled spliceosomes. To meet this high demand, snRNAs are often encoded by multiple gene copies, as is the case for tRNAs. Indeed, we found that mammals contain several minor spliceosome snRNA gene copies, particularly a high number of U6atac. Moreover, we found a higher number of U12 snRNA gene copies in carnivores compared to other mammals. Finally, there was a clear enrichment of U4atac snRNA gene copies in the primate lineage **(Fig. 5, Table S5)**.

**Figure 5.**
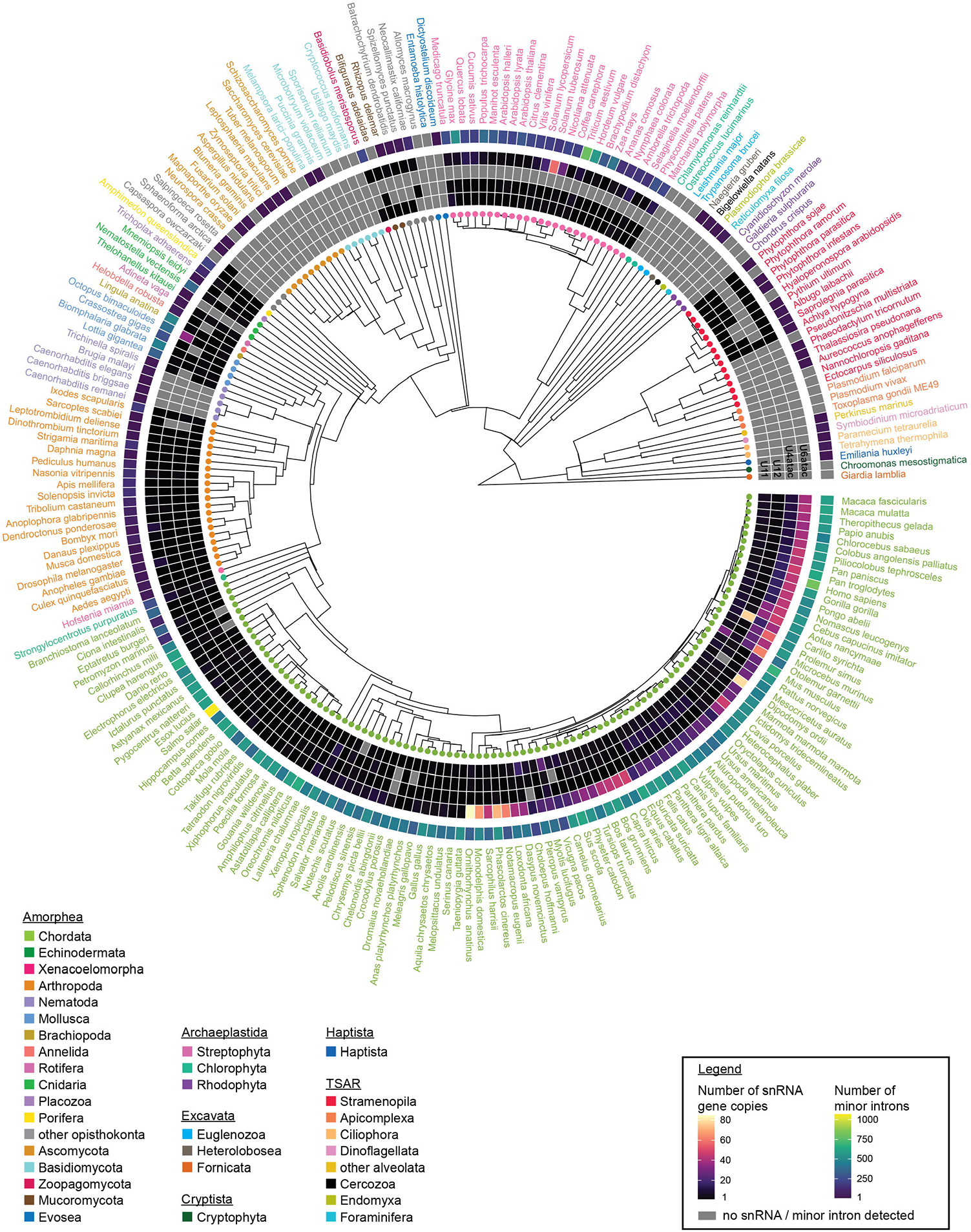
Detection of minor spliceosome-specific snRNAs in a diverse set of eukaryotic organisms. Circos plot with heatmap of the number of detected gene copies for U11, U12, U4atac and U6atac snRNA in a select set of eukaryotic organisms. The outer ring is a heatmap for the number of minor introns detected in each organism. Species are color-coded by phylum.

### Evolution of minor introns

The high conservation of minor introns and minor spliceosome components in disparate eukaryotic lineages supports the notion that minor introns are ancient. Nevertheless, the relatively high number of minor introns in chordates and mollusks, and the absence of minor introns and minor spliceosome components in many fungi, algae and protists suggests that minor introns are both lost and gained throughout evolution **(Fig. 4, 5)**. To gain more insight into the evolution of minor introns, we identified the orthologous introns of human minor introns across all investigated species **(Fig. 6)**. This revealed that most human minor introns were relatively conserved across chordates and mollusks. However, only few orthologous introns were also classified as minor introns in land plants, even though their genomes contain a comparable number of minor introns. Instead, human minor introns were orthologous to major-like/major introns in plants or were absent from the orthologous gene **(Fig. 6)**. We made similar observations for the orthologous introns in stramenopiles, which were either major introns or absent entirely **(Fig. 6)**. This suggests that these human minor introns have been lost from these lineages, through both conversion of intron class and homologous recombination with a reverse transcribed spliced mRNA transcript. The fact that plants and chordates have maintained a relatively high minor intron density suggests that their presence might provide a regulatory advantage in these lineages. In the past decades, several ideas have been postulated as to what this function of minor introns might be that ensured their conservation. Most prominently, it has been suggested that minor introns could regulate the expression of genes in which they are found (34, 35). Interestingly, we found that the most highly conserved minor introns were those found in the SLC9A genes, as well as the MAPK gene family. Given the important role of MAPK genes in the stress response, we therefore wondered if minor intron splicing might play a crucial role in this pathway, thereby explaining their high conservation.

**Figure 6.**
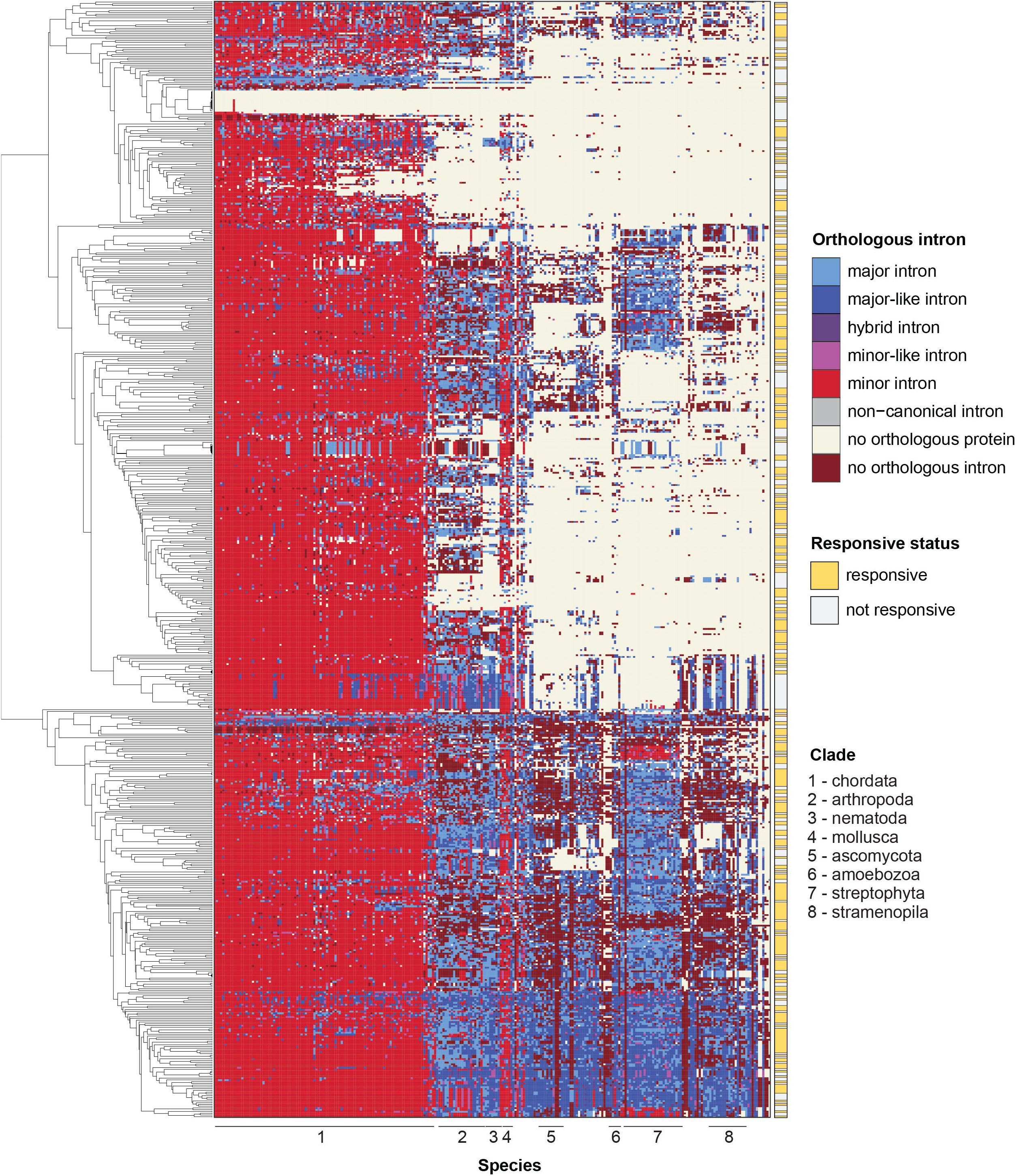
Orthology of human minor introns in a diverse set of eukaryotic organisms. Color-coded heatmap representing the identity of introns orthologous to human minor introns. Introns are clustered based on the identity of orthologous introns. Organisms are ordered by phylogeny, as in Figures 4 and 5.

### Minor intron splicing regulates gene expression in response to cold stress

Environmental stress, such as changes in temperature, is a constant selective pressure that organisms experience. Endotherms respond to environmental shifts in temperature by internally maintaining their core body temperature, while ectotherms such as fish and plants are susceptible to shifts in temperature. To test whether minor intron splicing is affected by environmental stresses such as cold exposure, we exposed the tropical poikilotherm Nile tilapia (*Oreochromis niloticus*) to acute cold stress by reducing the water temperature from 25°C to 14°C, at a rate of −1°C/h (36). RNAseq of the hypothalamus, a crucial thermoregulation center in the brain, revealed that over 20% of all minor introns were mis-spliced upon cold exposure, while only a small number of minor-like, hybrid and major-like introns were affected **(Fig. 7A-B)**. The MIGs with elevated minor intron retention significantly enriched for biological processes such as MAPK activity and transmembrane ion transport **(Fig. 7C)**. RT-PCR analysis of Nile tilapia exposed to chronic cold similarly showed elevated retention of minor introns **(Fig. 7D)**. These findings suggest that minor spliceosome, as reflected by elevated minor intron retention, responds to cold stress. Therefore, we extended our analysis to land plants, which are also exposed to a wide range of temperatures in nature. RNAseq analysis of cold exposed *Arabidopsis thaliana* and maize revealed that the splicing of minor introns was disproportionally affected, with 4 to 5-fold more minor introns retained than other intron types. In all, these findings place minor spliceosome activity downstream of cold stress, which results in elevated minor intron retention of a subset of MIGs involved in the cell stress response.

**Figure 7.**
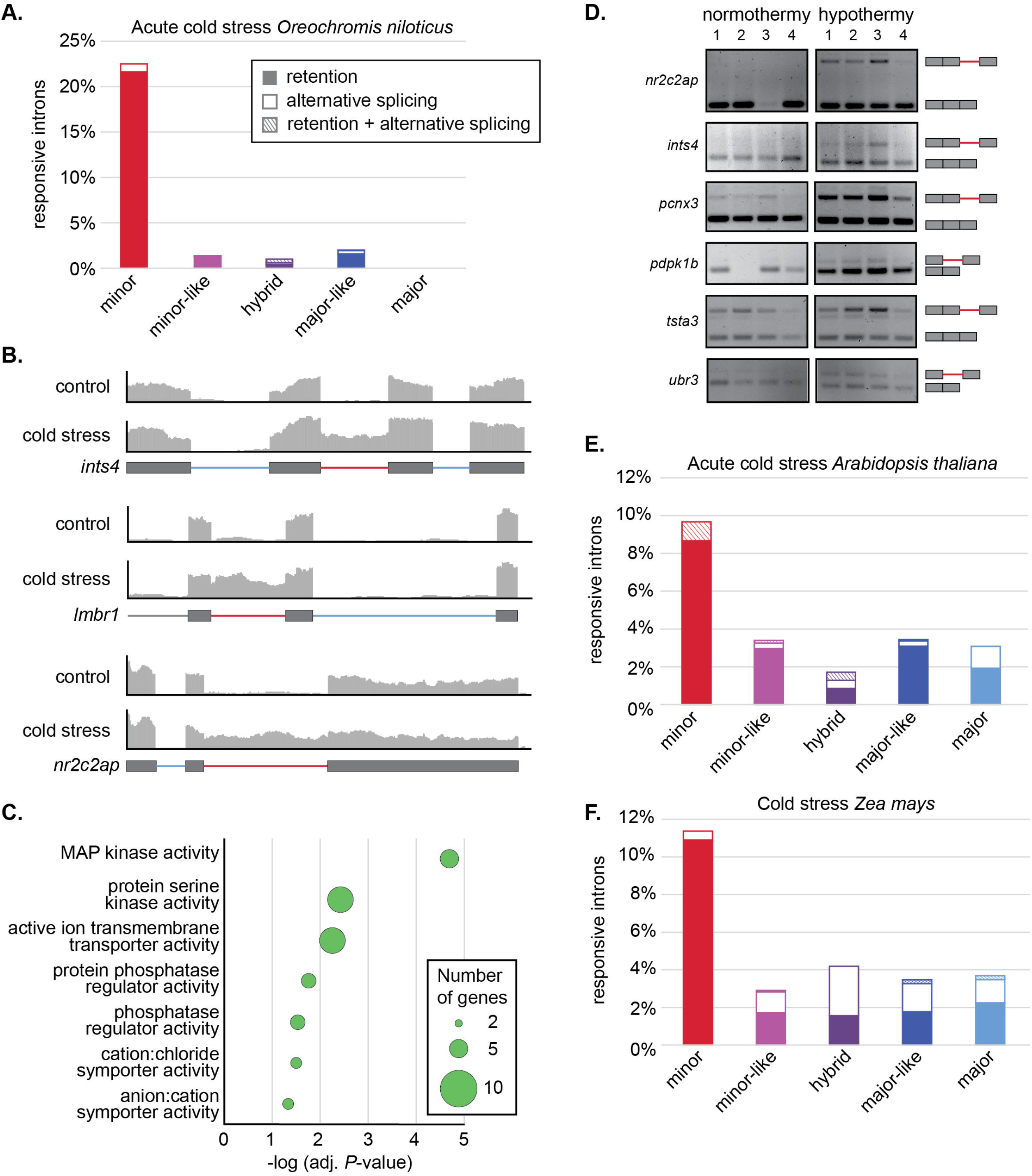
Cold stress in fish and plants results in elevated minor intron retention. **(A)** Bargraph showing the percentage of introns that are responsive to acute cold stress in Nile tilapia, either through elevated retention, alternative splicing or both. **(B)** Coverage plots for selected Nile tilapia genes whose splicing is affected by acute cold stress. Gene schematic with color-coded intron types are shown below. **(C)** Top hits from gene ontology analysis of the responsive minor introns in Nile tilapia. **(D)** RT-PCR analysis for select minor intron-containing genes showing elevated minor intron retention upon chronic cold stress in Nile tilapia (N=4 per condition). Gene schematics of Sanger-sequenced PCR products are shown on the right. **(E-F)** Bargraph showing the percentage of introns that are responsive to cold stress in **(E)** Arabidopsis

## Discussion

### Minor-like introns represent an intermediate in the conversion between minor and major introns

It is thought that an invasion of self-splicing group II introns into the genome was the source of major introns. Given that major introns are not randomly positioned in the genome but overrepresented in phase 0, suggests that these were not random insertion events. One model that has been proposed to explain this phase bias is that introns were inserted into proto-splice site sequences, such as AG|G. While these proto-splice sites are overrepresented in phase 0, this unequal distribution is not sufficient to fully explain the phase distribution of major introns (37, 38). Moreover, the proto-splice site hypothesis cannot explain the significant bias against phase 0 that is observed for minor introns **(Fig. S11)**. Instead, it has been suggested that the underrepresentation of phase 0 minor introns is the result of conversion of these introns to major introns (29). This might occur through the accumulation of sequential point mutations of key nucleotides in the splice sites of minor introns. Indeed, it has been shown that three point mutations in the splice sites of the minor intron of P120 are sufficient for the major spliceosome to be able to splice this intron (20). Additionally, identification of orthologous intron pairs in distant species, where one is major and the other minor, has previously been used as support for the idea that minor introns can be converted to major introns (18, 39). To gain insight into how these transitions take place during evolution, we employed a more refined intron classification strategy to capture intermediate intron states. We found that minor-like introns are present in almost all eukaryotic lineages and propose that they represent an intermediate in the conversion of minor to major introns **(Fig. 4)**. In support of this idea, most minor-like introns are found in orthologous intron clusters with significant enrichment of minor introns **(Fig. 3D)**. While the intronic sequence elements of minor-like introns are very similar to that of minor introns, the presence of a guanine at the −1 position of the 5’SS and +1 position of the 3’SS are a feature of major introns **(Fig. 1C)**. The −1G nucleotide is one of the most conserved nucleotides in major introns and is not only used for base-paring with U1 snRNA, but also with U6 and U5 snRNA (40). Moreover, the disruption of splicing due to mutations in the 5’SS of major introns can be rescued by converting the −1 nucleotide to a G, underscoring the importance of this nucleotide in 5’SS recognition and major intron splicing. It has been proposed that this guanine plays a role in stabilizing the pre-catalytic major spliceosome by Watson-Crick base-pairing (40). This would open the possibility that minor-like introns possessing a guanine at the −1 position can be spliced by the major spliceosome. Our data revealed that the splicing of minor-like introns lacking this −1G was affected by minor spliceosome inhibition, while the splicing of minor-like introns with a guanine at this position was unaffected **(Fig. 3A)**. This finding suggests that the highly abundant major spliceosome normally competes for the splicing of minor-like introns with the less abundant minor spliceosome (41). The mere presence of this guanine at some minor-like introns might stabilize the pre-catalytic major spliceosome sufficiently, such that the minor spliceosome is not necessary. On the other hand, absence of this nucleotide does not leave enough Watson-Crick interactions between the minor-like splice sites with the major spliceosome snRNA, so that the splicing of these introns depends on the minor spliceosome. The bias against guanines at the −1 and +1 position of the exon/intron boundary can explain why responsive minor-like introns were often found in phase 1 and 2, as these nucleotides are part of the proto-splice site sequence overrepresented in phase 0 **(Fig. 3C)**. In all, these findings suggest that minor-like introns represent a natural snapshot capturing the conversion of minor introns into major introns.

### The possibility of a hybrid spliceosome

The energetic burden associated with maintaining a spliceosome has likely led to an evolutionary pressure to streamline its composition. Indeed, mass spectrometry analysis of the major spliceosome B and C complexes in human and yeast has revealed that yeast spliceosomes contain drastically fewer proteins than those in humans (42). Attempts at bioinformatically identifying orthologs of these ∼60 core spliceosome proteins in more extant species such as *Giardia lamblia* and *Encephalitozoon cuniculi*, revealed the existence of only 30-35 proteins (43, 44). Further evidence of the dramatic reduction of spliceosome size in ancient species stems from *Cyanidioschyzon merolae*, which lacks both U1 snRNA and all associated proteins (45). Despite the absence of many spliceosome components, introns in these species are spliced, thereby underscoring the extreme diversity and flexibility that exists in the splicing reaction. Here, we have detected minor introns in the genomes of several species that lack specific minor spliceosome snRNAs **(Fig. S14)**. Most notably, these include the land plants, water mould gyrista and several fungal species of the mucoromycote and zoopagomycote lineage, for which we identified gene copies for U11, U12 and U6atac snRNA, but not U4atac **(Fig. 5; Fig. S1)**. The absence of U4atac snRNA can mean one of three things: 1) the sequence and structure has diverged so significantly that they escaped detection by our bioinformatics pipeline, 2) minor splicing in these species can occur without U4atac, and/or 3) U4atac can be replaced by U4 snRNA in these species.

While we cannot formally exclude the first two options, several lines of evidence lead us to favor the third option. First, even though U4atac snRNA is the least conserved of all snRNAs **(Fig. 5)**, mutations in *RNU4ATAC* have been linked to several congenital developmental diseases (46–49). This suggests that its importance in the minor intron splicing reaction might in fact be restricted to humans and/or vertebrates. Indeed, we observed a noticeable increase in the number of genes encoding U4atac snRNA in primates **(Fig. 5)**, suggesting that U4atac snRNA is particularly important for this lineage. Second, unlike U11, U12 and U6atac snRNA, which bind to proteins unique to the minor spliceosome, U4atac snRNA has no such protein interactions reported. While a few U4atac/U6atac-specific proteins have been identified, they are all found in the mono-U6atac snRNP fraction and/or bound to U6atac snRNA in structure (50, 51). Thus, it is easy to imagine that U4 snRNA could replace U4atac snRNA, so long as it can base-pair with U6atac snRNA. Indeed, biochemical experiments have already revealed that the introduction of compensatory point mutations in U4 snRNA that allow for its base-pairing with U6atac snRNA is sufficient to activate minor intron splicing (52). Together, this opens the exciting possibility that non-metazoan species lacking U4atac snRNA can splice minor introns using a hybrid spliceosome that consists of both major and minor spliceosome components.

Invoking a hybrid machinery is not too far-fetched, considering that the minor spliceosome already uses many major spliceosome proteins, along with U5 snRNA. In fact, the existence of such a hybrid spliceosome has been proposed for *Giardia lamblia*, which contains snRNAs that possess features of both major and minor spliceosome snRNAs. For instance, the U6 snRNA of *G. lamblia* is truncated at the 5’ end compared to the human U6 snRNA and contains a 3’ stem loop that is characteristic of the human U6atac (53). This stem loop is important for minor intron splicing, as creation of a chimeric U6atac consisting of the human snRNA with the 3’ stem loop of *Arabidopsis thaliana, Drosophila melanogaster, Trichinella spiralis*, or *Phytophtora infestans* could only partially, or not at all, support the splicing of a minor intron in the P120 gene reporter (54). The existence of a hybrid spliceosome could also explain how hybrid introns, which possess sequence elements of both major and minor introns, are spliced **(Fig. 1C)**. In all, this interchangeable aspect of spliceosome components could also explain how many minor introns are still spliced normally when the minor spliceosome is inhibited. While we show the existence of many minor and minor-like introns that are highly dependent on the proper expression and function of all minor spliceosome components, the fact that many minor and minor-like introns remain unresponsive to minor spliceosome inhibition argues for the presence of a hybrid spliceosome.

### Regulatory advantage of minor introns in stress response

As discussed earlier, maintenance of one spliceosome poses a high energetic burden, thereby raising the question as to why an organism would maintain two spliceosomes. Indeed, several lineages have lost minor introns and the minor spliceosome altogether **(Fig. 4-5)**. Nevertheless, the relatively high conservation of minor introns in vertebrates and land plants suggests that perhaps they provide an evolutionary advantage in these lineages. Several models have been developed to explain the function of minor introns. For instance, it has been postulated that minor introns are spliced more slowly than neighboring major introns in the same gene, thereby creating a bottleneck in the expression of these MIGs (34). Indeed, at steady-state, many minor introns are retained at higher levels than major introns, and intron-retained transcripts are stuck in the nucleus (55). Minor intron splicing efficiency can be increased through stabilization of U6atac snRNA, a rate-limiting component of the minor spliceosome with high turnover kinetics (35). Thus, increasing U6atac half-life might provide a rapid means to effectively regulate minor intron splicing and MIG expression in conditions of cellular stress. The overrepresentation of MIGs in certain biological processes has been proposed as another reason for the maintenance of minor introns. For example, MIGs are enriched in the essentialome and have been shown to be important for cell cycle regulation (56). However, because cell cycle regulation is an essential process for any organism, both unicellular and multicellular, the high conservation of minor introns specifically in animals and plants suggests that there is another biological function of MIGs that can provide evolutionary advantages.

Here we found that the most highly conserved minor introns are positioned in genes of the MAPK family **(Fig. 6)**, which play a central role in the signal transduction of diverse stress responses. Moreover, we found that minor introns are highly conserved in the ubiquitously expressed SLC9A (NHE) branch of the Na+/H+ exchangers that regulate organellar pH (57). The expression and activity of these receptors has been shown to play an important role in conferring resistance to salt and osmotic stress in Arabidopsis (58, 59). Finally, minor introns in genes that play a role in glycosylation are highly conserved in plants and animals **(Fig. 6)** (60). Glycosylation has recently been shown to be important in regulating the cellular stress response, and specifically the oxidative stress response in plants and cancer cells (61, 62). Based on these findings, we wondered whether minor introns might have been maintained in vertebrates and plants due to their capacity to respond to environmental stresses, such as osmotic, oxidative, heat and cold stress. In support of this hypothesis, we observed widespread minor intron retention in the hypothalamus of *Oreochromis niloticus* exposed to both acute and chronic cold stress, as well as in cold-exposed *Arabidopsis thaliana* and *Zea mays* **(Fig. 7)**. While cold stress has previously been shown to result in alternative splicing and changes in gene expression (36, 63, 64), our data indicates that it disproportionally affects the splicing of minor introns and expression of MIGs. The mechanism by which this specificity is achieved remains elusive and warrants further research. Nevertheless, MIGs whose splicing is affected by cold stress are involved in biological pathways regulating energy metabolism and internal ion concentration. These data suggest that ectotherms like fish and plants have adopted the minor spliceosome as part of their stress response mechanisms, thereby providing a rationale for the paradoxical high conservation of minor introns in animals and plants.

In all, our intron classification strategy has provided insight into the evolution of minor introns through the identification of minor-like introns. We have integrated this data into the Minor Intron Database (midb.pnb.uconn.edu) and hope that this resource will aid in expanding the study of splicing in non-model organisms. Moreover, we believe that the creation of a comprehensive list of the introns whose splicing is affected by minor spliceosome inhibition will aid in the understanding of the molecular pathogenesis of spliceosomopathies.

## Data availability

The intron classification for all 263 genomes can be found on the Minor Intron Database (MIDB), and can be accessed at midb.pnb.uconn.edu.

The transcriptomic datasets generated and analyzed during this study are available in the Gene Expression Omnibus. Accession numbers for individual datasets can be found in **Table S2**.

## Funding

Funding for this study was obtained from the National Institute of Neurological Disorders and Stroke [R01NS102538 and R21NS096684 to RNK].

### Conflict of interest statement

None declared.

## Acknowledgements

We would like to acknowledge Dr. Ion Mandoiu from the Computer Science Engineering Department of the University of Connecticut for creating the necessary infrastructure to perform all bioinformatics analysis. We would to acknowledge the generosity of Dr. Mazoyer from the Centre de Recherche en Neurosciences de Lyon, who shared the transcriptomic data obtained from MOPD1 patients (13). Moreover, we would like to thank Emma Schwoerer and Nick Mcintosh for providing helpful feedback on the design and construction of MIDB. Finally, we would like to thank Kaitlin Girardini for her feedback on the manuscript.

## Author contributions

Conceptualization - AMO and RNK; methodology – AMO, CFS and RNK; software – AMO, CFS, AW; investigation – AMO, CFS, IA, KD, SM, JKH, AC, JB and RNK; resources – JKH, JB, RNK; writing – AMO and RNK; supervision - RNK; funding acquisition - RNK.

## Notes

http://midb.pnb.uconn.edu

### Competing Interest Statement

The authors have declared no competing interest.

