## Supplemental Figures for "Taxonomy of introns, their evolution, and the role of minor introns in stress response"

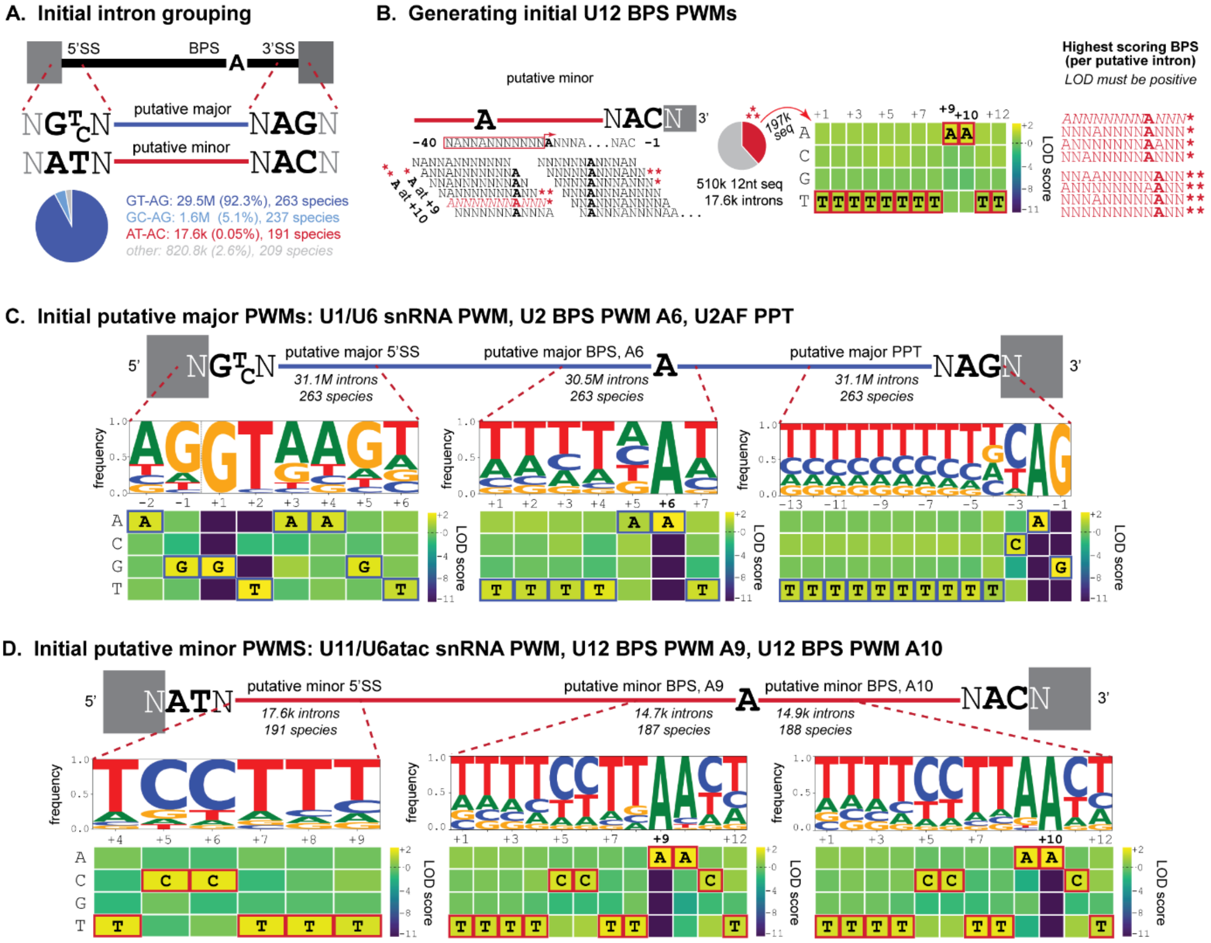


**Figure S1. Generation of initial PWMs. (A)** Introns were grouped based on terminal dinucleotides. **(B)** Schematic displaying the sliding window utilized to create an initial minor-type BPS PWM. **(C-D)** Frequency logos and LOD scores of initial (C) major and (D) minor PWMs.

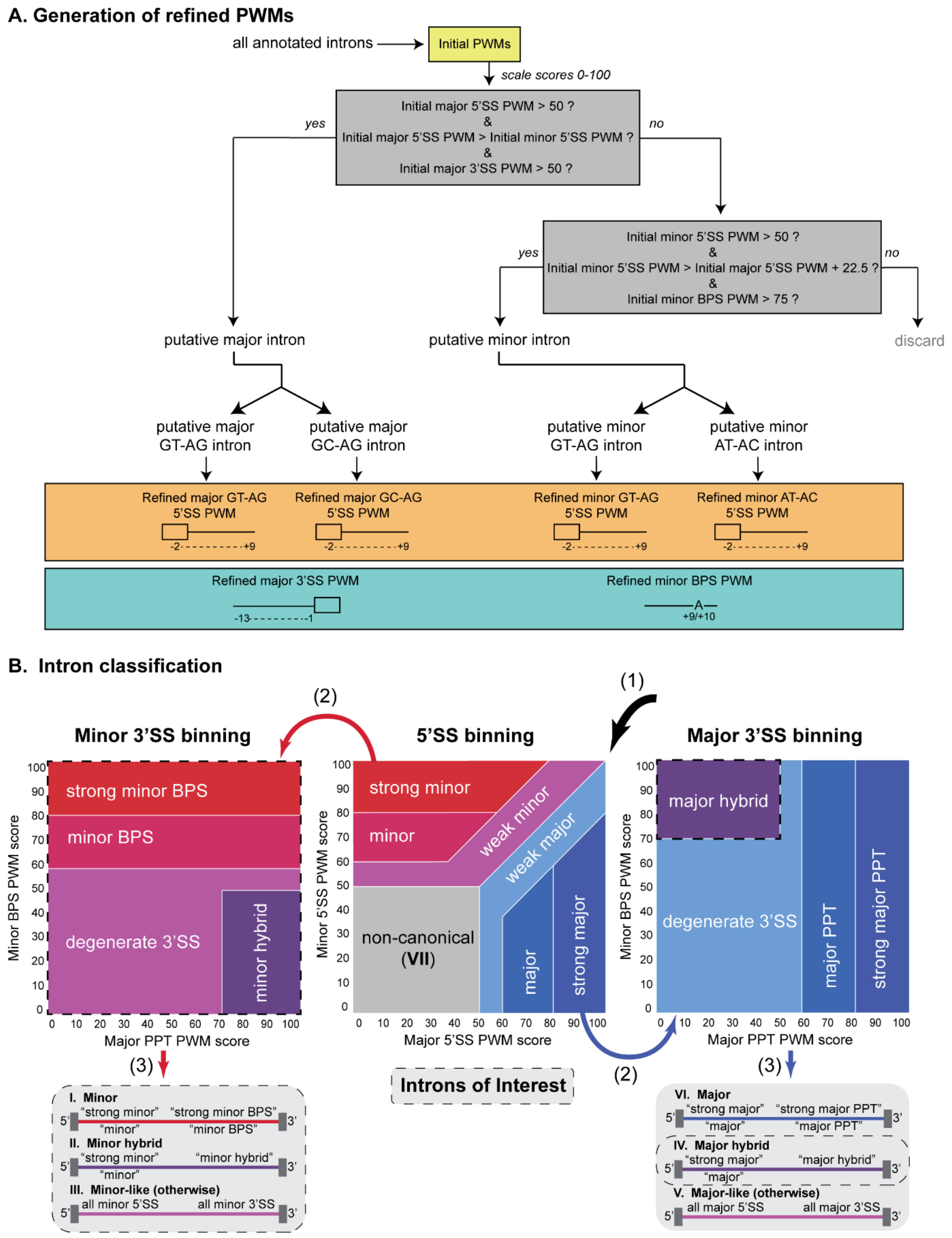


**Figure S2. Generation of refined PWMs and classification of introns. (A)** Flowchart depicting the generation of refined PWMs used for intron scoring. **(B)** Classification of introns based on scores for the 5'SS, BPS and PPT PWM. Introns are first binned based on 5'SS (1), and depending on their classification further scored using the major or minor 3'SS classification scheme (2). Introns with different subclassifications are then merged into one of 7 intron categories (3): (I) minor, (II) minor-hybrid, (III) minor-like, (IV) major-hybrid, (V) major-like, (VI) major, (VII) non-canonical. The major-hybrid and minor-hybrid categories are merged for majority of the analyses.

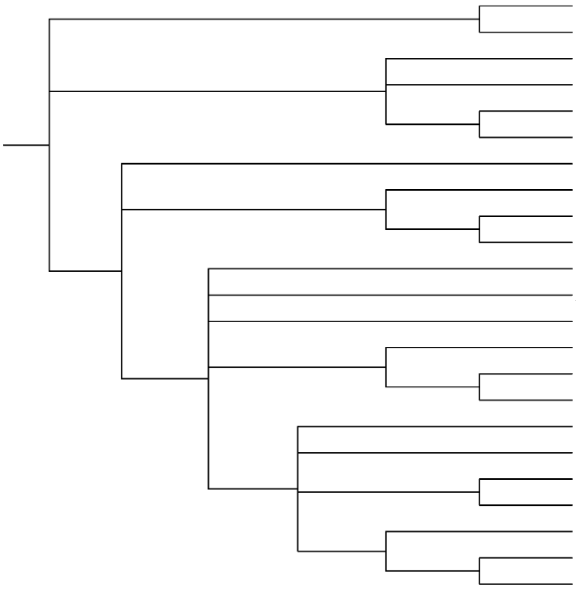

| Species | MIDB | IAOD | U12DB | ERISdb |
|---|---|---|---|---|
| *Saccharomyces cerevisiae* | 0 | 0 | 0 | |
| *Schizosaccharomyces pombe* | 0 | 0 | | |
| *Glycine max* | 553 | 509 | | 654 |
| *Arabidopsis thaliana* | 300 | 270 | 246 | 409 |
| *Oryza sativa Japonica* | 251 | 282 | | 481 |
| *Zea mays* | 422 | 250 | | 783 |
| *Caenorhabditis elegans* | 6 | 0 | 0 | |
| *Apis mellifera* | 132 | 128 | 34 | |
| *Drosophila melanogaster* | 16 | 17 | 16 | |
| *Anopheles gambiae* | 23 | 25 | 4 | |
| *Gallus gallus* | 546 | 508 | 399 | |
| *Xenopus tropicalis* | 568 | 416 | 329 | |
| *Ciona intestinalis* | 88 | 100 | 116 | |
| *Danio rerio* | 684 | 662 | 298 | |
| *Tetraodon nigroviridis* | 381 | 460 | 309 | |
| *Takifugu rubripes* | 586 | 483 | 337 | |
| *Bos taurus* | 578 | 592 | 496 | |
| *Canis lupus familiaris* | 610 | 559 | 558 | |
| *Rattus norvegicus* | 571 | 602 | 524 | |
| *Mus musculus* | 707 | 657 | 555 | |
| *Macaca mulatta* | 637 | 521 | 479 | |
| *Homo sapiens* | 850 | 681 | 695 | |
| *Pan troglodytes* | 657 | 623 | 524 | |

**Figure S3. Comparison of minor intron counts obtained by different databases.**

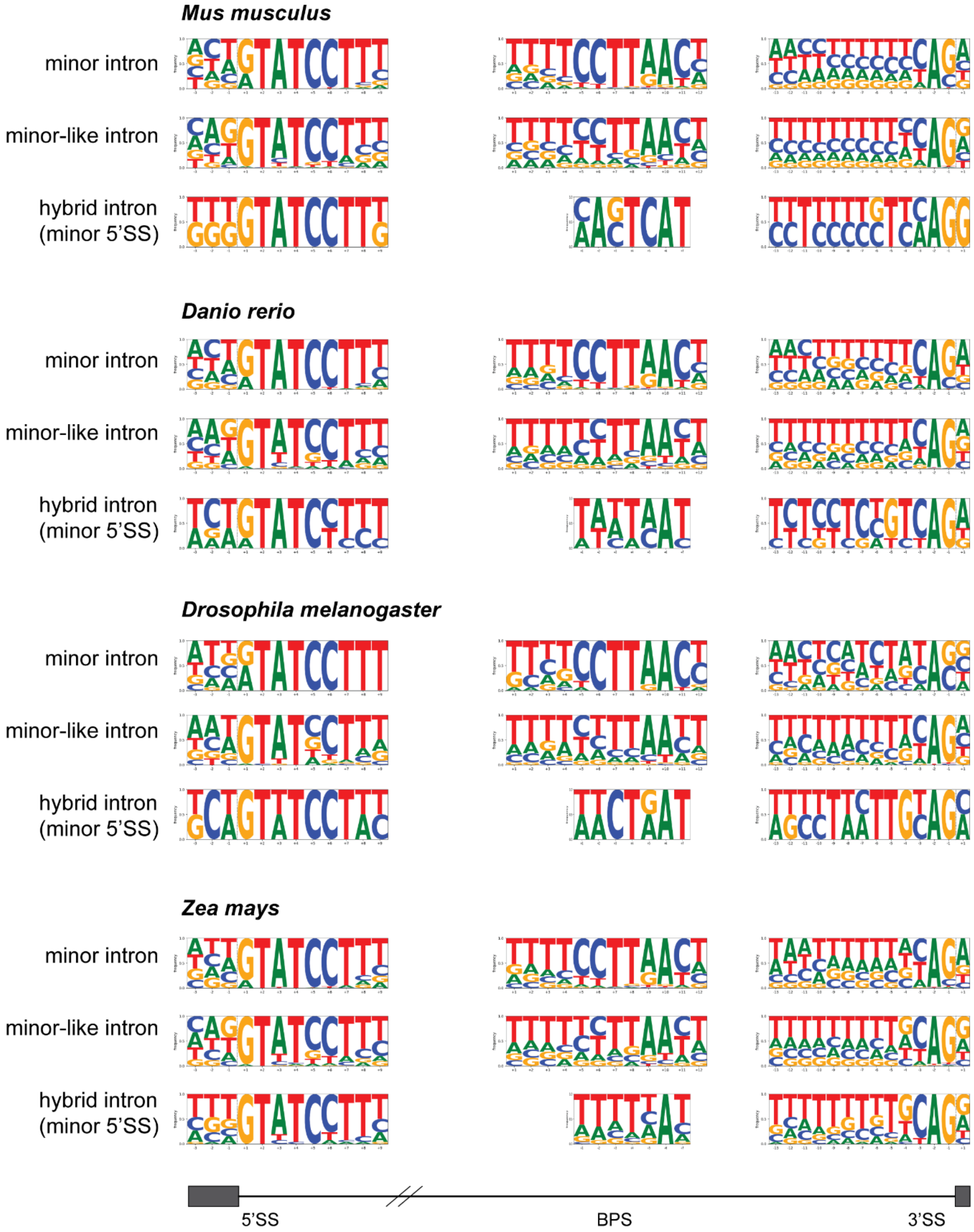


**Figure S4. Consensus sequences of minor, minor-like and minor-hybrid introns at the 5'SS, BPS and 3'SS.**

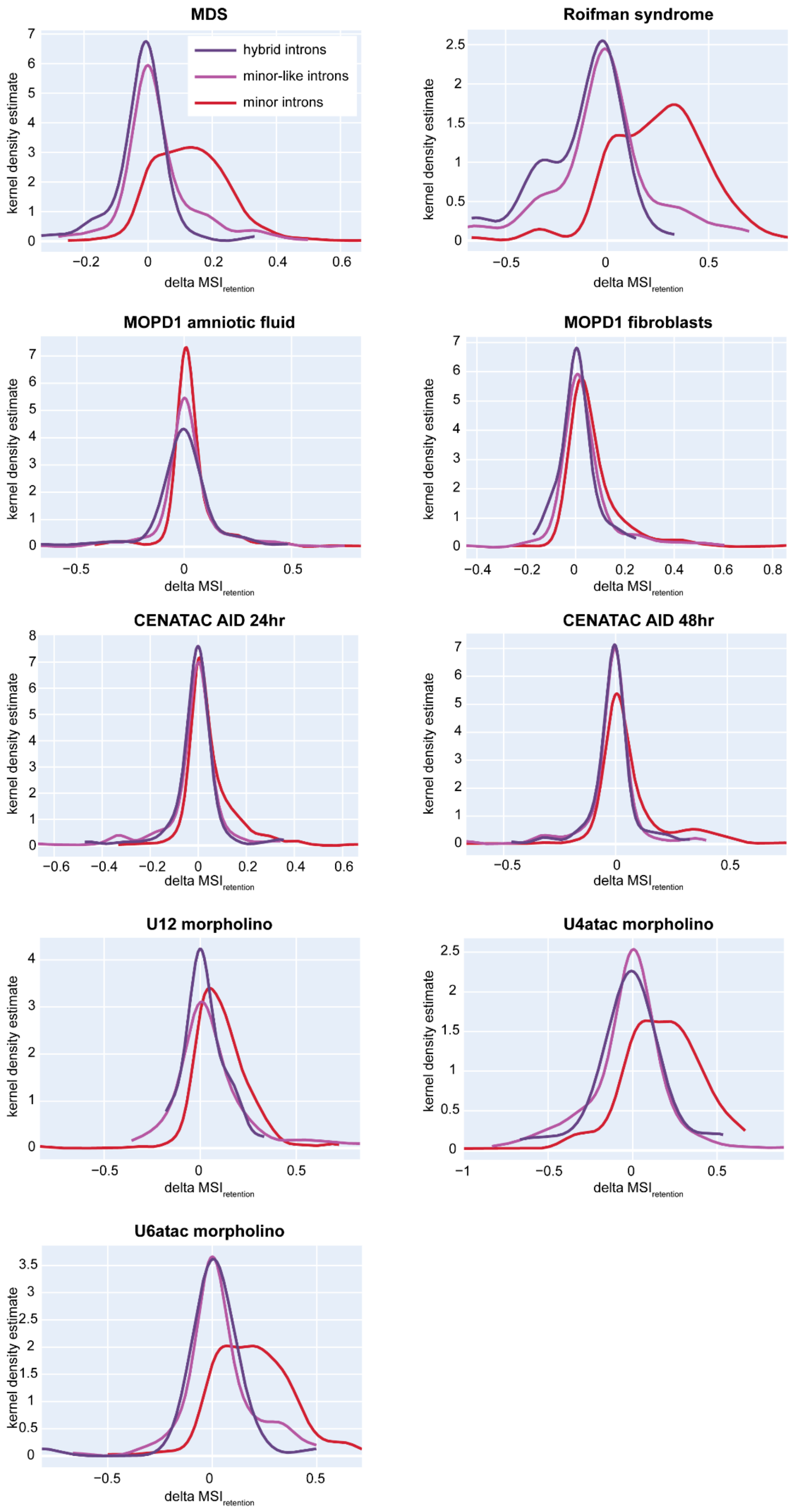


**Figure S5. Minor intron splicing defects in human datasets with minor spliceosome inhibition.** Density plot for the level of retention of minor, minor-like and hybrid introns in the experimental compared to respective control condition. MSI=mis-splicing index.

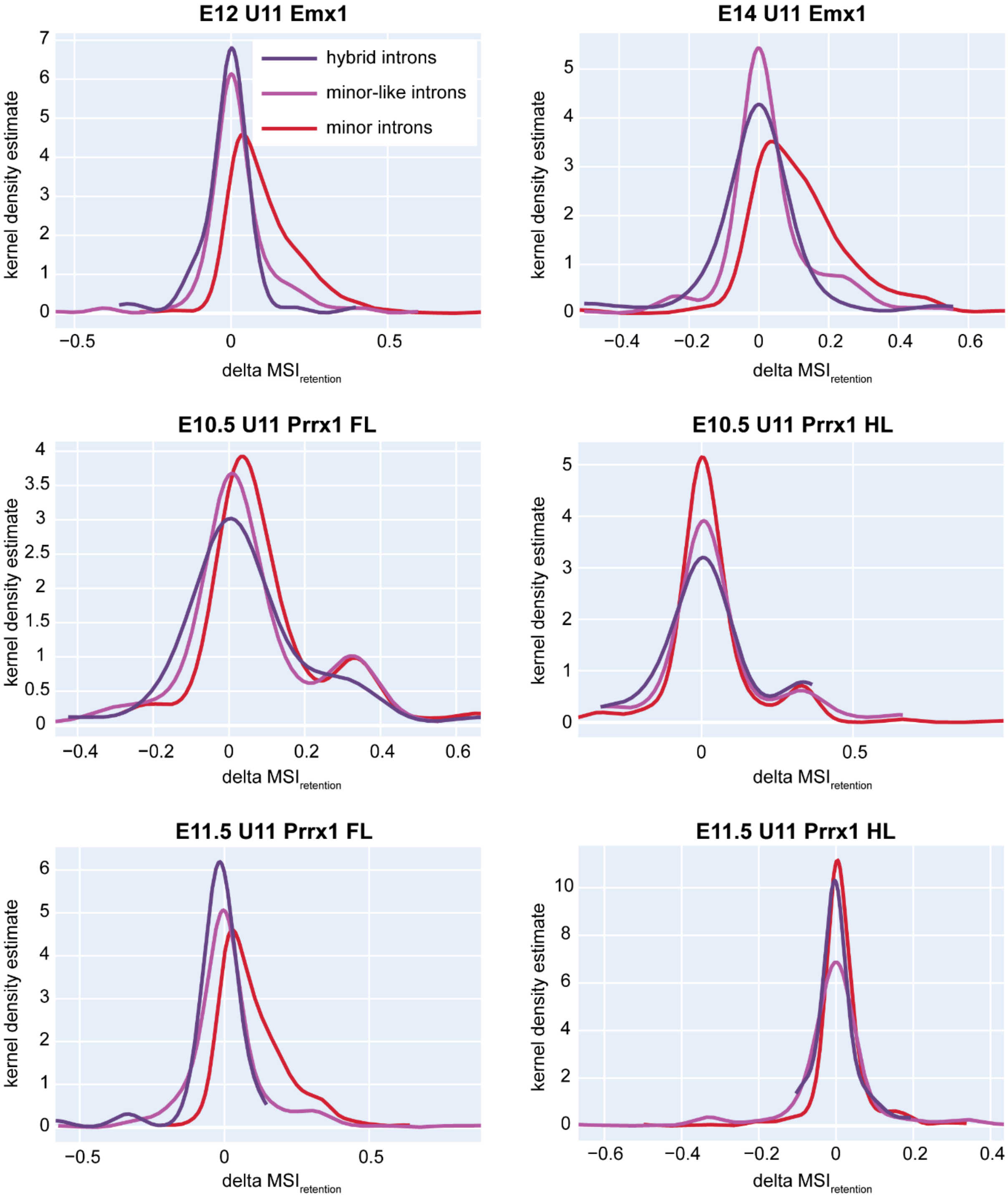


**Figure S6. Minor intron splicing defects in mouse datasets with minor spliceosome inhibition.** Density plot for the level of retention of minor, minor-like and hybrid introns in the experimental compared to respective control condition. MSI=mis-splicing index.

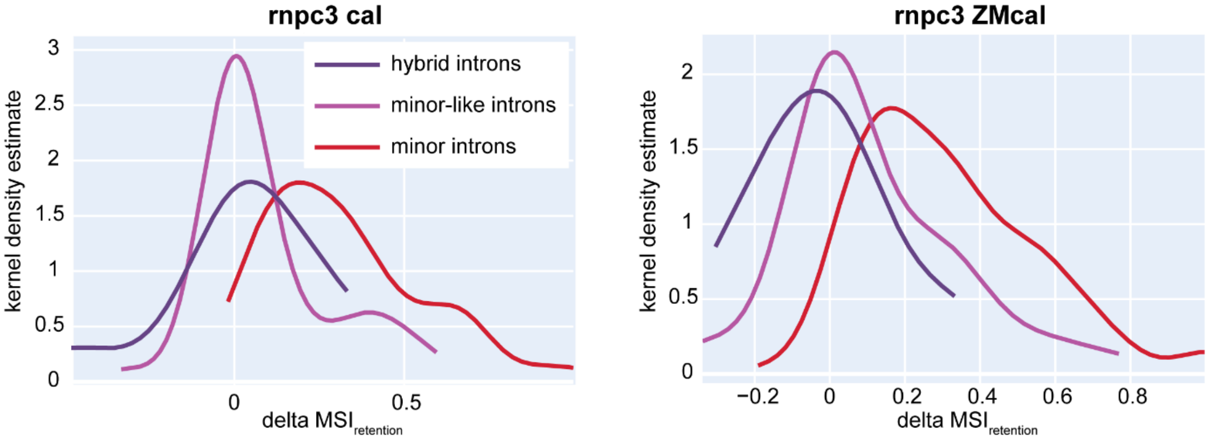


**Figure S7. Minor intron splicing defects in zebrafish datasets with minor spliceosome inhibition.** Density plot for the level of retention of minor, minor-like and hybrid introns in the experimental compared to respective control condition. MSI=mis-splicing index.

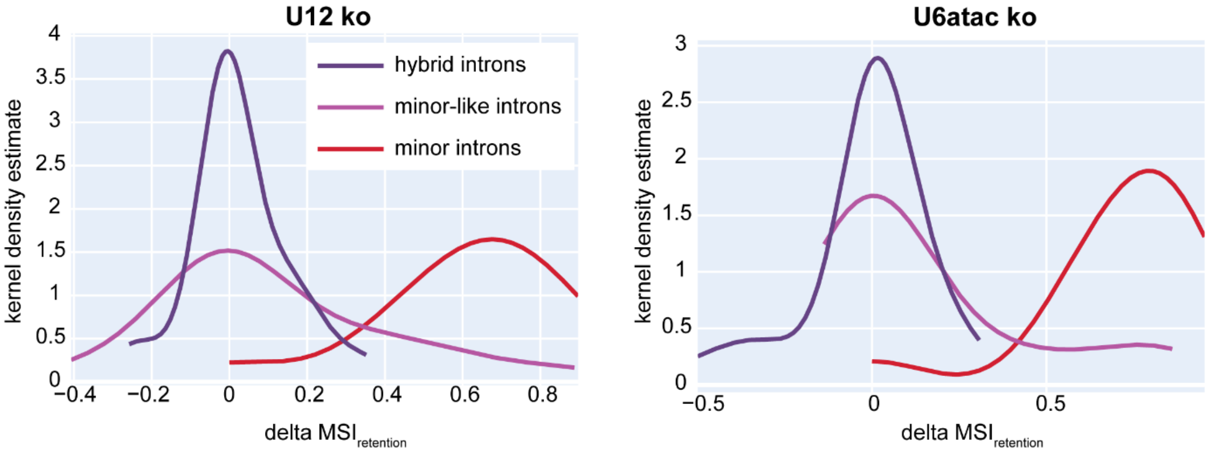


**Figure S8. Minor intron splicing defects in fruit fly datasets with minor spliceosome inhibition.** Density plot for the level of retention of minor, minor-like and hybrid introns in the experimental compared to respective control condition. MSI=mis-splicing index.

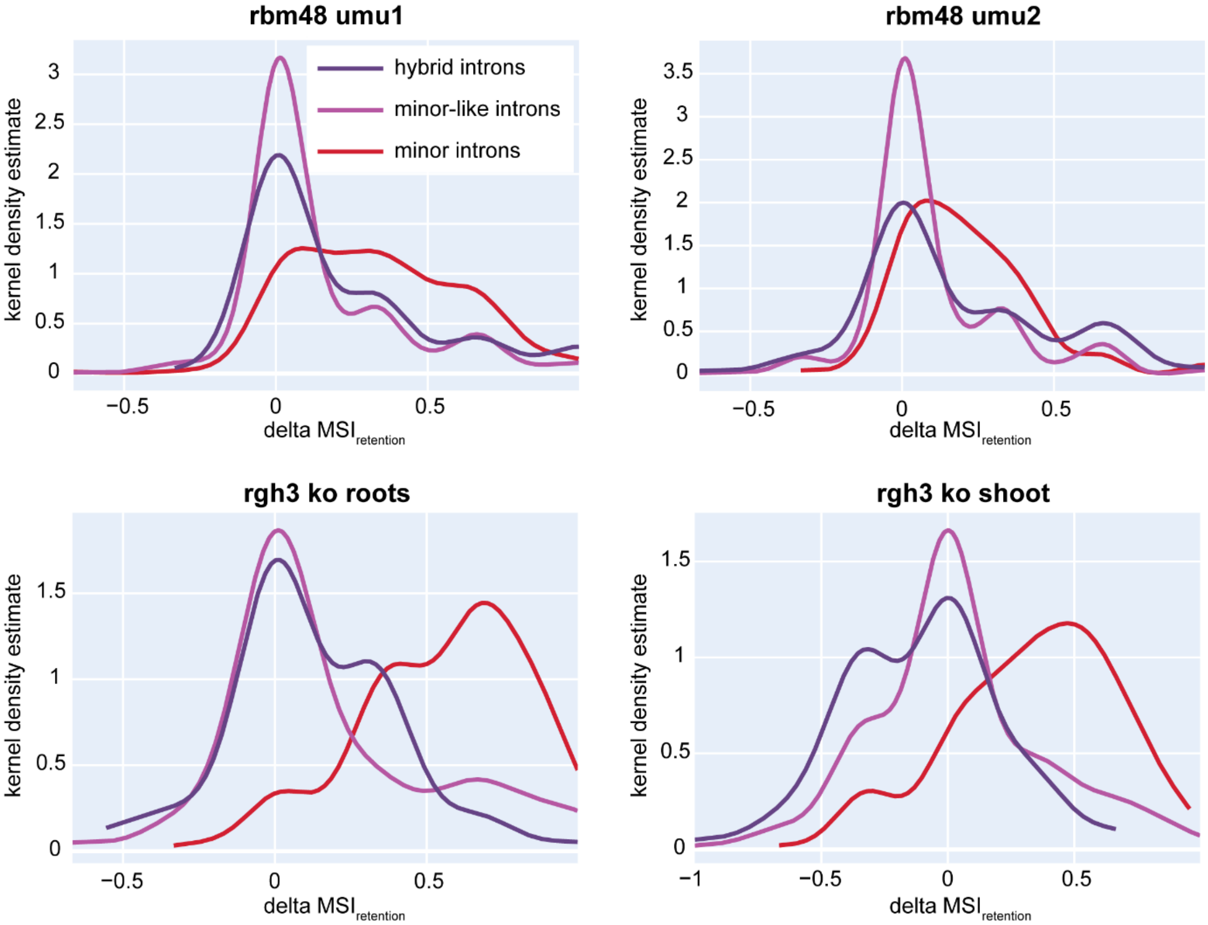


**Figure S9. Minor intron splicing defects in maize datasets with minor spliceosome inhibition.** Density plot for the level of retention of minor, minor-like and hybrid introns in the experimental compared to respective control condition. MSI=mis-splicing index.

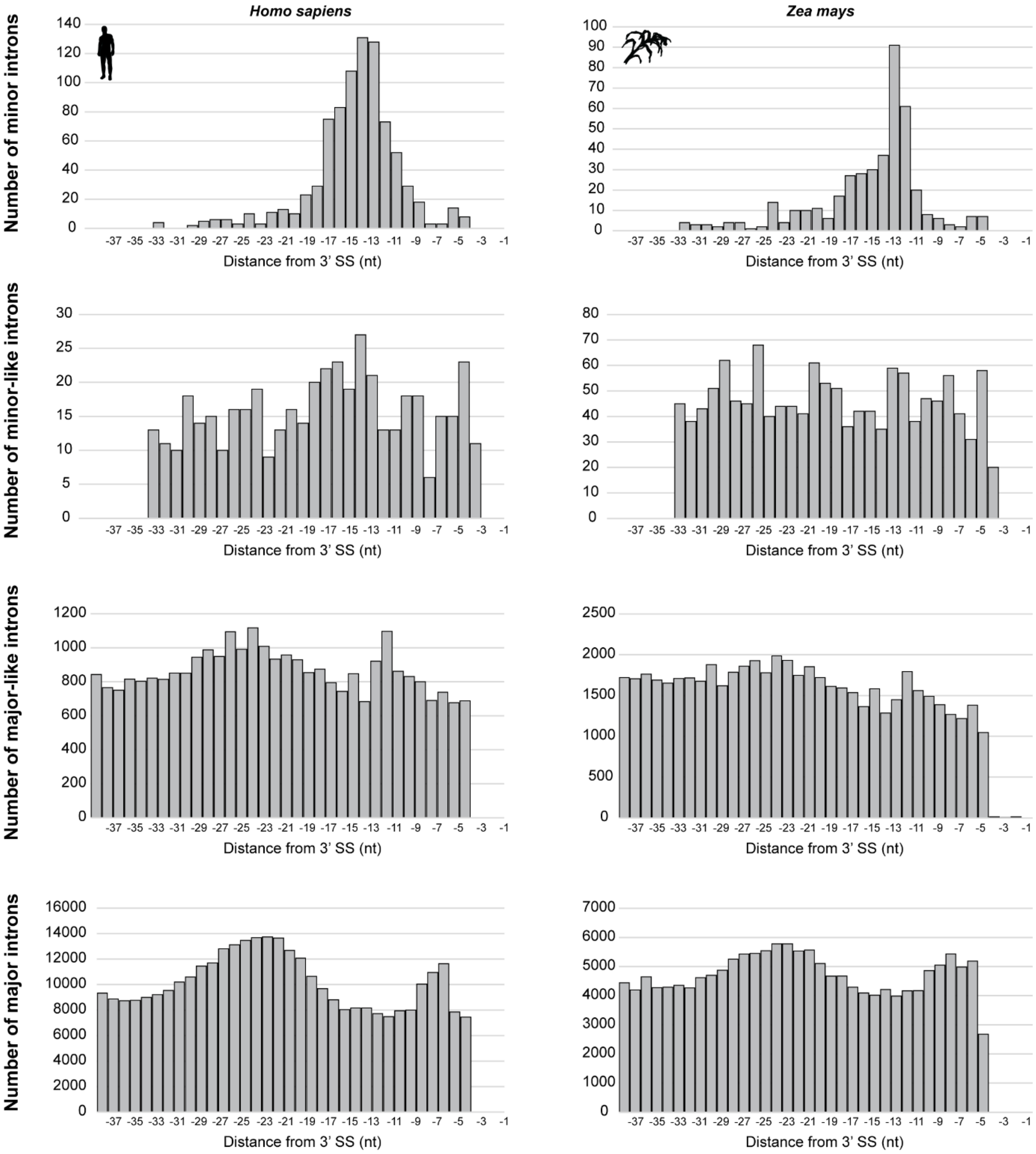


**Figure S10. Distance of branchpoint adenosine from the 3' exon-intron boundary for different intron classes in human and maize.**

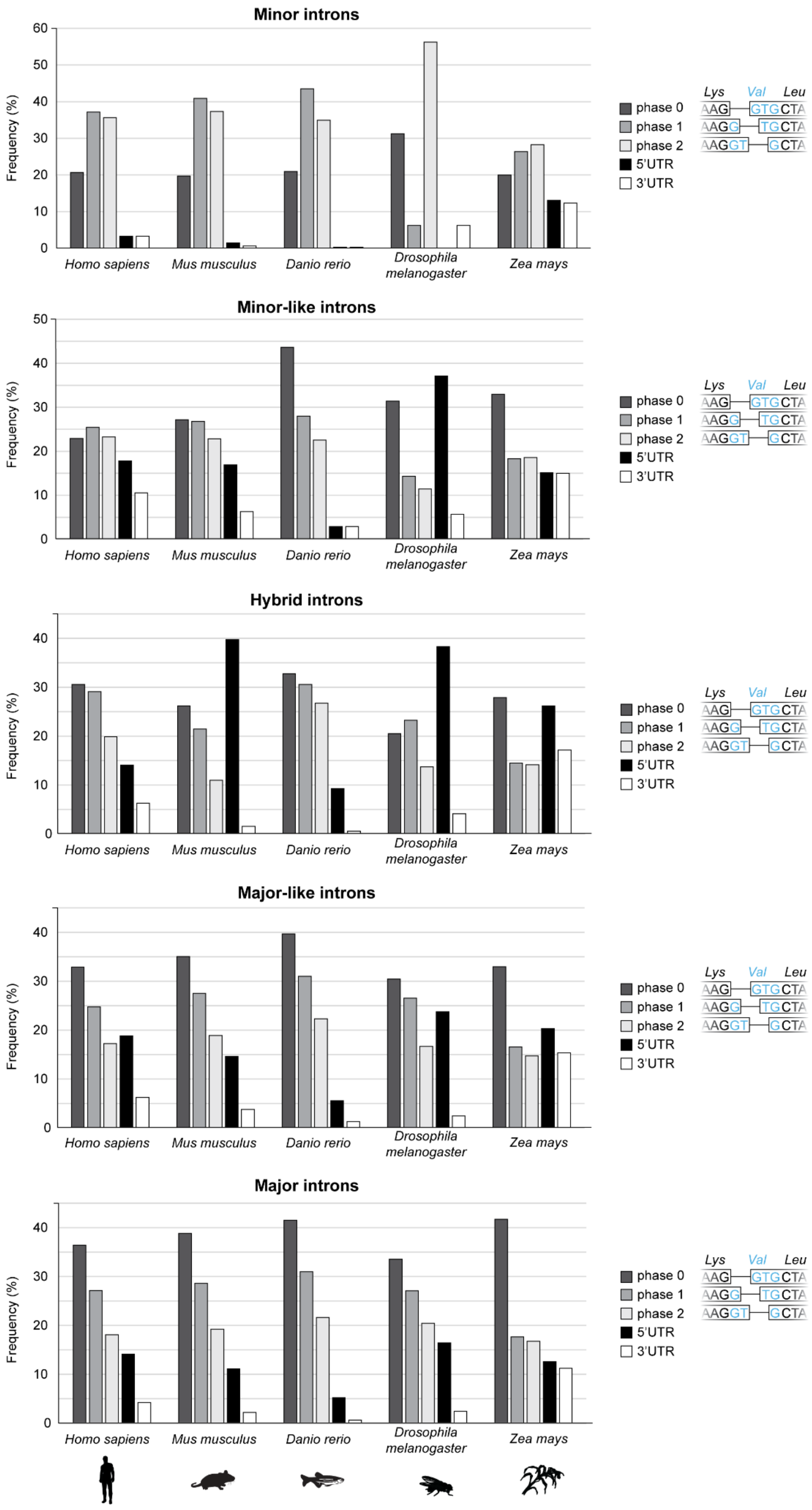


**Figure S11. Phase distribution of different intron types in model organisms.**

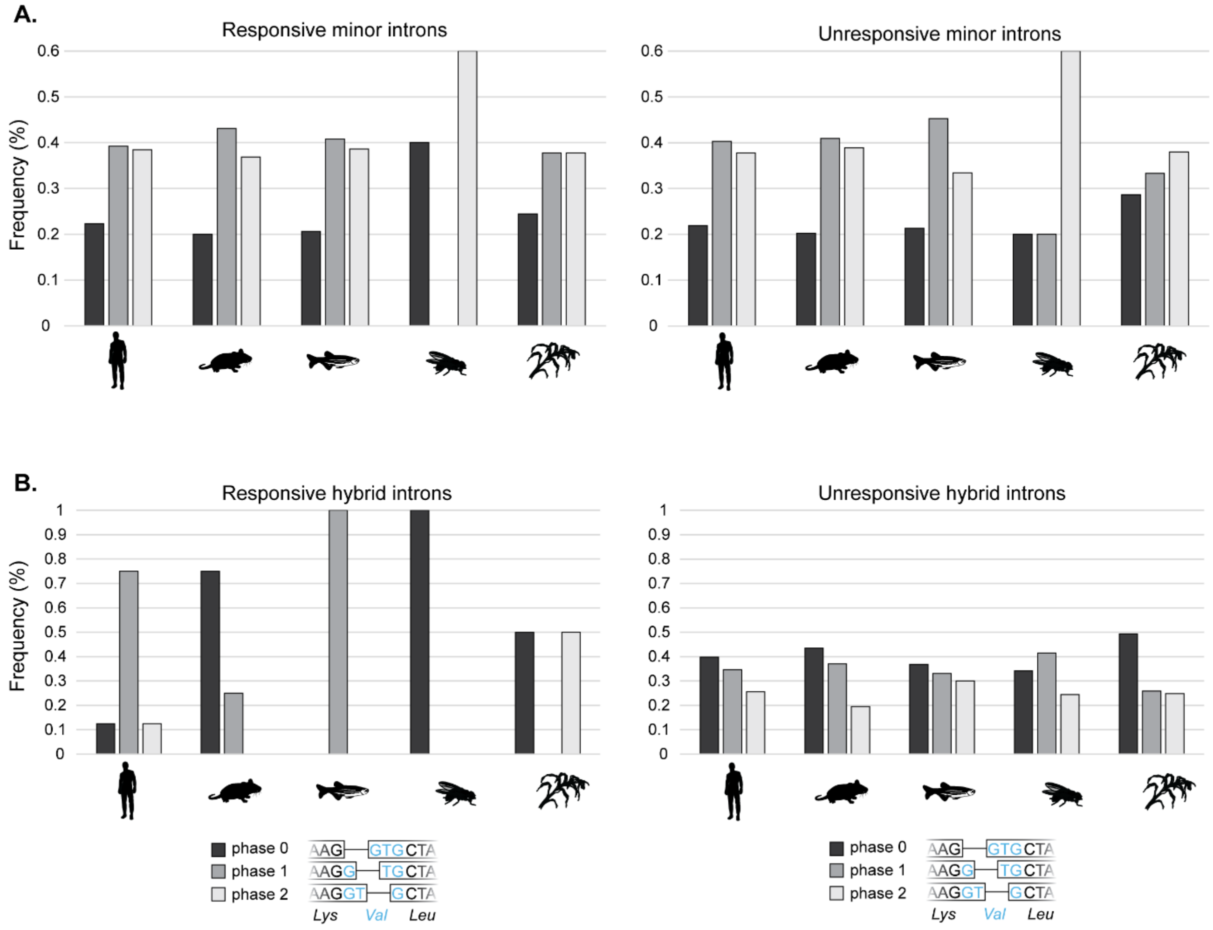


**Figure S12. Phase distribution of minor and hybrid introns by responsive status. (A-B)** Phase distribution of responsive and unresponsive **(A)** minor and **(B)** hybrid introns in model organisms.

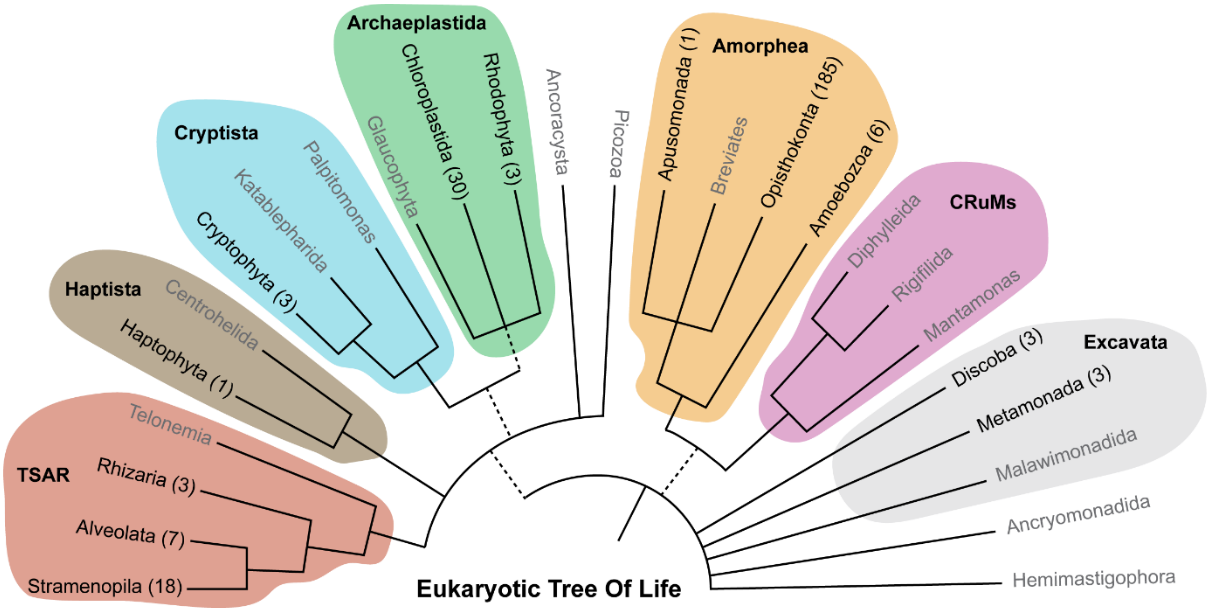


**Figure S13. The eukaryotic tree of life.** Number analyzed genomes for intron classification are listed for each clade between brackets.

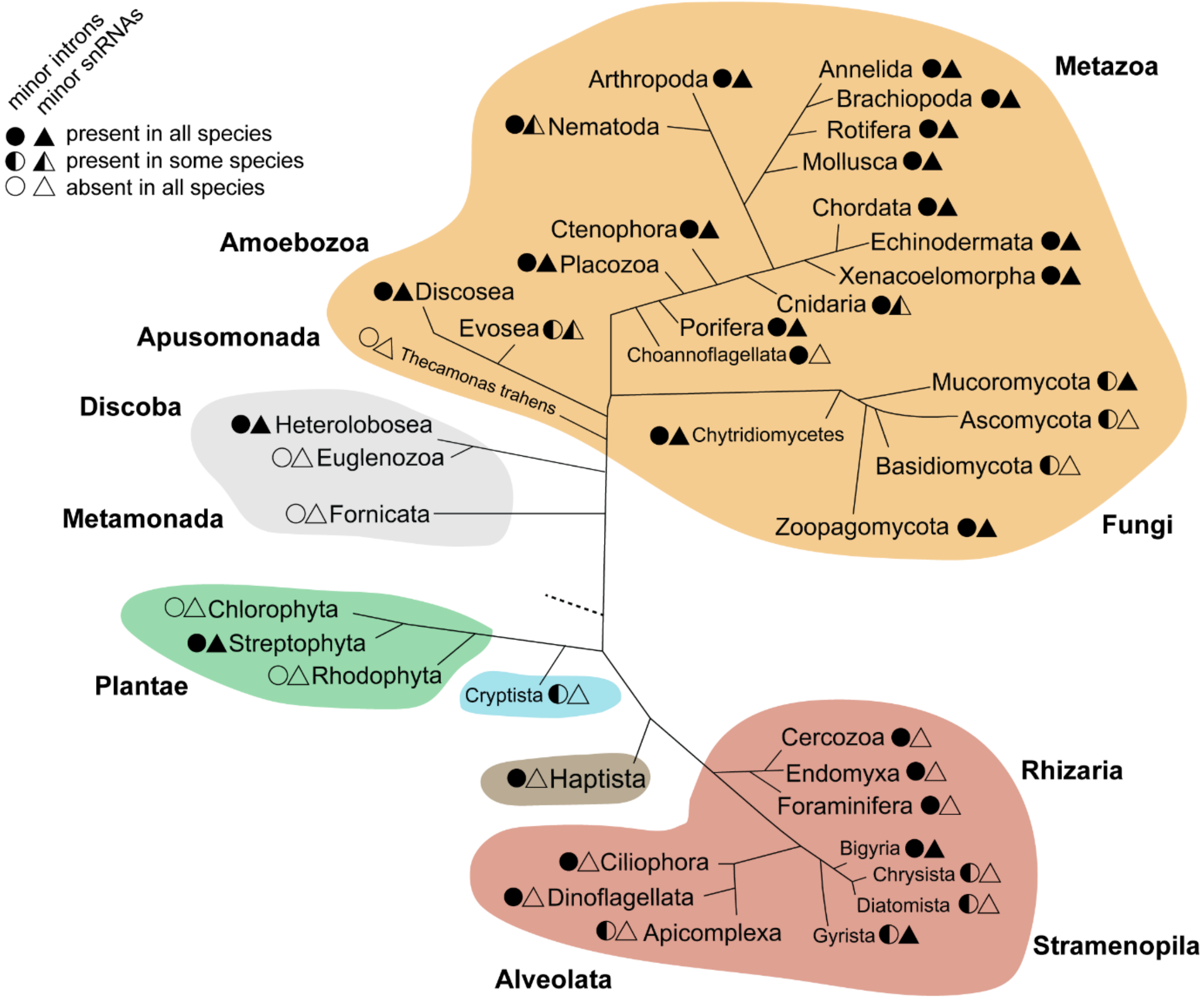


**Figure S14. Minor intron splicing across the eukaryotic tree of life.** Presence of minor introns (circles) and minor spliceosome snRNAs (triangles) are listed for each clade.
